# Immune-like glycan-sensing and horizontally-acquired glycan-processing orchestrate host control in a microbial endosymbiosis

**DOI:** 10.1101/2024.09.14.613017

**Authors:** Benjamin H. Jenkins, Estelle S. Kilias, Fiona R. Savory, Megan E. S. Sørensen, Camille Poirier, Victoria Attah, Georgia C. Drew, Luis J. Galindo, Guy Leonard, Duncan D. Cameron, Michael A. Brockhurst, David S. Milner, Thomas A. Richards

## Abstract

Endosymbiosis was a key factor in the evolution of eukaryotic cellular complexity. Yet the mechanisms that allow host regulation of intracellular symbionts, a pre-requisite for stable endosymbiosis and subsequent organelle evolution, are largely unknown. Here, we describe an immune-like glycan-sensing/processing network, partly assembled through horizontal gene-transfers (HGTs), that enables *Paramecium bursaria* to control its algal endosymbionts. Using phylogenetics, RNA-interference (RNAi), and metabolite exposure experiments, we show that *P. bursaria* regulates endosymbiont destruction using glycan-sensing/processing – a system that includes a eukaryotic-wide chitin-binding chitinase-like protein (*CLP*) localized to the host phago-lysosome. RNAi of *CLP* alters expression of eight glycan-processing genes, including two prokaryote-derived HGTs, during endosymbiont destruction. Furthermore, glycan-sensing/processing dynamically regulates endosymbiont number in *P. bursaria*, plasticity crucial to maximize host fitness across ecological conditions. *CLP* is homologous to a human phagocyte-associated innate immune factor, revealing how immune functions can be alternatively adapted and expanded, partly through HGT, enabling endosymbiotic control.

**Graphical Abstract:** 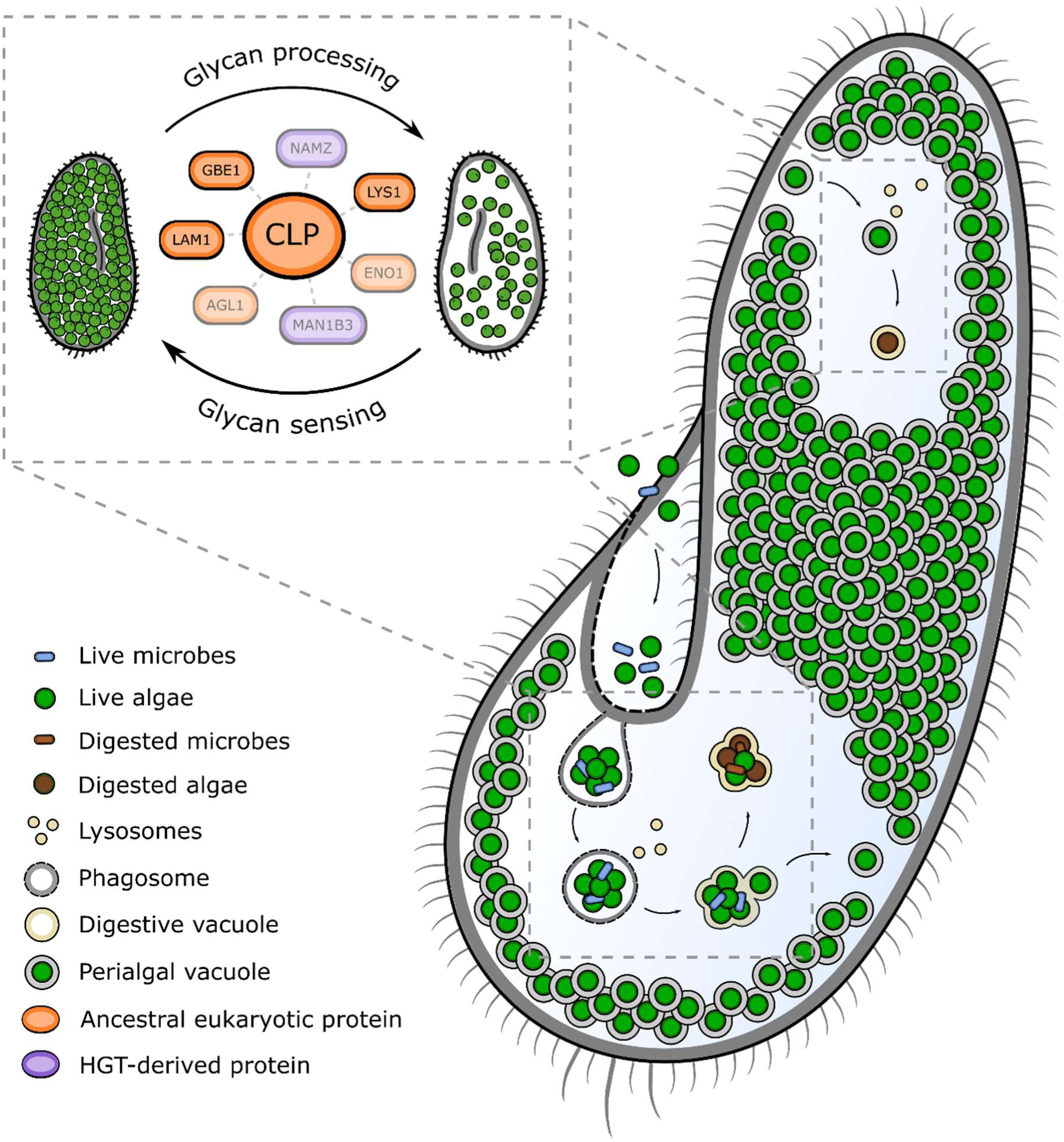

---

Endosymbiosis played a crucial role in the evolution of eukaryotic cellular complexity^1–4^ and underpins diverse ecologically and medically-significant host-microbe interactions (e.g., ^5–11^). Long-term maintenance of a stable endosymbiosis requires a network of host mechanisms that regulate endosymbiont functions, allowing the host to maximize the fitness benefit of the interaction under fluctuating ecological conditions^12–15^. Orchestrating this network requires extensive genetic and cellular re-modelling by the host; however, the function and evolution of these mechanisms are poorly understood.

The ciliate protist, *Paramecium bursaria* (*Pb*), is an RNAi-tractable emerging model organism amenable for studying facultative endosymbiotic interactions^16–19^. Each cell maintains hundreds of intracellular green algae from the *Chlorella*-clade (e.g., *Chlorella* & *Micractinium*^20^) in a symbiosis involving photosynthesis-derived metabolite exchange^21–24^. Endosymbiotic algae are housed in modified host phagosomes (perialgal vacuoles) localized beneath the host cell cortex by microtubules^25–27^. *Pb* can regulate endosymbiont load through detachment and lysosomal fusion of the perialgal vacuole to trigger endosymbiont digestion^26–29^. In this manner, *Pb* dynamically controls the endosymbiont population in response to factors such as light intensity to maximize host fitness, often at the expense of individual endosymbiont fitness^13,29–31^. *Chlorella* algae possess cell walls composed of complex glycans (predominantly oligomannosides and chitin^32–34^) which a host must break-down to digest and thereby control its endosymbionts. Accordingly, the presence of *Chlorella* endosymbionts leads to increased expression of host chitin-processing genes in both *Pb*^35^ and *Hydra viridis*^36^, a metazoan host which also forms symbioses with *Chlorella* species^36^.

Glycan processing and sensing requires functionally diverse glycosyl-hydrolases (GHs)^37,38^. GHs are widespread but discontinuously distributed in eukaryotes, with these enzymes performing important structural, nutritional and signaling functions in many ecologically-significant interactions^39–42^. This includes break-down and sensing of glycans derived from hosts, pathogens, other types of symbionts, or food^35,36,39,42–45^. Chitin oligosaccharides, such as the monosaccharide N-acetyl-glucosamine (GlcNAc), are common cues of innate immunity that can trigger both pathogen defense and symbiotic maintenance^42,44–48^. The spread of GHs in eukaryotes has been further facilitated by horizontal gene transfer (HGT), particularly among pathogens of glycan-producing hosts^49–56^.

Here, we investigate the glycan-sensing and processing network in *Pb*, focusing on a GlcNAc oligosaccharide binding chitinase-like protein (*CLP*) which is homologous to an innate immune-factor protein in humans. We demonstrate using RNAi that host *CLP* perturbation can disrupt endosymbiont destruction and alters the expression of intracellular signaling systems and GH genes, suggesting a role in host sensing cascades and down-stream glycan-processing. We demonstrate that 9 of the 37 GH genes in *Pb* (24%) were acquired via prokaryote-to-eukaryote HGT, including two (a lysozyme – *NAMZ*; and a mannosidase – MAN1B3) with altered expression during *CLP* RNAi perturbation, indicating how HGT can augment ancestral functions of preexisting genes in order to facilitate a host-microbe interaction. We show that RNAi perturbation of an additional GH gene (a eukaryotic lysozyme – *LYS1*) also alters host control of endosymbiotic algae. We propose that a functional network of immune-like glycan sensing, dependent upon *CLP*, and augmented by HGT-derived glycan processing genes, has been repurposed as a mechanism of endosymbiont control in *Pb* through the regulation of endosymbiont destruction. We discuss how such networks represent a key innovation enabling hosts to regulate microbial endosymbionts.

## Results

### Horizontal gene-transfer has shaped glycan-processing in P. bursaria

Host-microbe interactions are often governed by glycan-processing^44–48^. To identify the repertoire of glycan-processing genes in *Pb* 186b^19^, we designed a bioinformatic annotation pipeline for the discovery of carbohydrate-active enzymes^37^ (https://github.com/benjaminhjenkins/CAZyme_survey). This identified a total of 37 putative glycosyl-hydrolase (GH) (25 GH families; **Fig. S1a**) and 12 putative glycosyl-transferase (GT) (10 GT families; **Fig. S1b**) encoding genes. Comparative analyses revealed a mosaic distribution of GH and GT genes across the ciliates, including multiple instances of gene duplication and loss (**Supplementary Dataset S1**).

Phylogenetic analyses were then conducted to identify candidate horizontal gene-transfers (HGT) among the *Pb* GH genes. These resolved tree topologies for nine of the 37 *Pb* GH genes (24.3%) that were best explained by HGT^57^ from within the prokaryotes into a eukaryotic ancestor lineage of *Pb*, given current genome sampling (**Fig. 1** – four HGT events involving five *Pb* genes, discussed further below; see **Supplementary Table S1** and **Fig. S2** for details on the additional four HGTs identified). Four of the putative HGT-derived genes were shown to be acquired specifically in the *Pb* stem lineage (**Fig. 1a/c** & **S2 b/c**) and were present in all five *Pb* strains assessed, ruling out genome project contamination. Comparison of trimmed mean M (TMM) values of transcript abundance in *Pb* 186b^19^ (under scramble control RNAi conditions, see below) demonstrates transcription of each putative HGT-derived gene with the exception of *MAN1B2*, indicating possible pseudogenization of *MAN1B2* following *MAN1B1/2* gene duplication. Each putative HGT-derived gene contained introns (**Supplementary Table S1**), including short introns characteristic of *Paramecia*^19^. Furthermore, each proposed open reading frame (ORF) contained between 9 and 43 UAG/UAA codons (representing 2.8% to 5.6% of codons per ORF; **Supplementary Table S1**), which in *Paramecium*^58^ code for glutamine rather than stop codons as translated in the universal genetic code used by the prokaryotic progenitor lineage of the HGT. Collectively, these data suggest that the putative HGT-derived genes have been integrated to function within the host eukaryotic genome.

**Figure 1:**
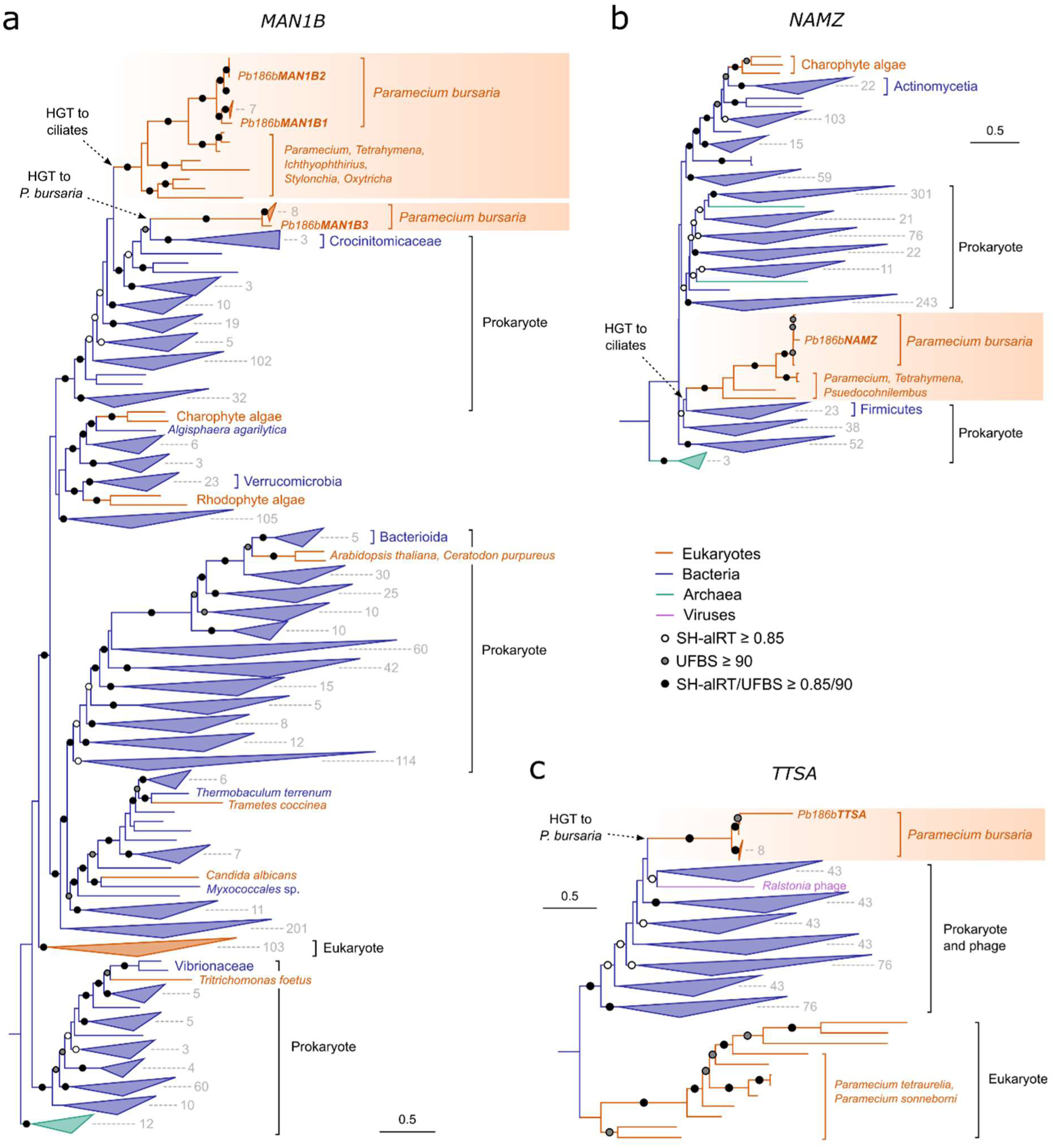
Horizontal gene-transfer has shaped glycan-processing in *P. bursaria*. Maximum likelihood phylogenies of *Pb β-mannosidase* (*MAN1B*; ***a***), *NamZ-like lysozyme* (*NAMZ*; ***b***), and *TtsA-like lysozyme* (*TTSA*; ***c***) genes. Topologies of these trees provide strong support for prokaryote-to-eukaryote HGT, with two independent incidences of HGT and a subsequent gene duplication of *MAN1B1/2* predicted to have occurred in *Pb* (***a***). We observed no evidence of *MAN1B2* transcription (**Supplementary Table S1**) indicating possible pseudogenization of *MAN1B2* in *Pb* 186b. Trees were generated in IQ-TREE using Q.pfam+R10 (***a***), Q.pfam+I+R10 (***b***), or Q.pfam+R7 (***c***) substitution models selected by Model Finder, and statistical support assessed using SH-aLRT and ultra-fast bootstraps (UFBS; *n =* 1000). This figure shows four cases of HGT of GH gene families, for phylogenies of four additional predicted HGT-derived GH encoding genes in *Pb* see **Fig. S2.**

The proposed HGT-derived genes were annotated as encoding putative *β-mannosidase* (*MAN1B1-3* and *MANEB*; **Fig. 1a** and **Fig. S2d**), *lysozyme* (*NAMZ* and *TTSA*; **Fig. 1b/c**), *β-glucosidase* (GAB; **Fig. S2a**), *cellulase* (CELA1; **Fig. S2b**), and *β-fructofuranosidase* (INV; **Fig. S2c**) enzymes. Interestingly, the proposed functions of these HGT-derived genes are enriched for cleavage of mannose (*MAN1B1-3*, *MANEB*) and GlcNAc (*NAMZ*, *TTSA*) containing substrates – components of both bacterial food and the endosymbiotic algal cell wall^32–34^ – hinting that these HGT-acquired functions are important for mixotrophic functions and endosymbiont control. These data reveal a mosaic distribution of glycan-processing genes in *Pb* and the broader ciliate group that has been augmented by prokaryote-to-eukaryote HGT.

### RNAi of a subset of glycan-processing genes perturbs host-mediated endosymbiont destruction, while perturbation of CLP determines expression of glycan-processing genes

Next, we sought to understand whether *Pb* GH genes function in endosymbiont control. We conducted an RNA-interference (RNAi) screen on *Pb* populations grown in microwell plates and treated with cycloheximide to induce endosymbiont destruction^28,29^. Per-host fluorescence (a proxy for endosymbiotic algal load^59^) was measured on an ImageXpress Pico automated cell imaging system, and compared to ‘non-hit’ RNAi control (scramble) populations to assess the effect of GH transcript perturbation on endosymbiont destruction (**Fig. 2a**). RNAi of *u2af* –a splicing factor in *Pb* required for host growth– was used as a positive RNAi control^16^. The RNAi screen was conducted twice (two biological replicates per treatment with six technical replicate populations each; *n* = 12) and mean per-host fluorescence data were assessed with a mixed effects model to account for uncontrolled sources of variance. This revealed that 22% of observable variance could be attributed to RNAi-based gene perturbation, 24% could be attributed to random effects (biological replicate and plate positional artifacts), and 54% remained unexplained. Due to the high unexplained variability and considerable influence of confounding factors, we have been conservative in our interpretation of these data and consider the results set out below as trends.

**Figure 2:**
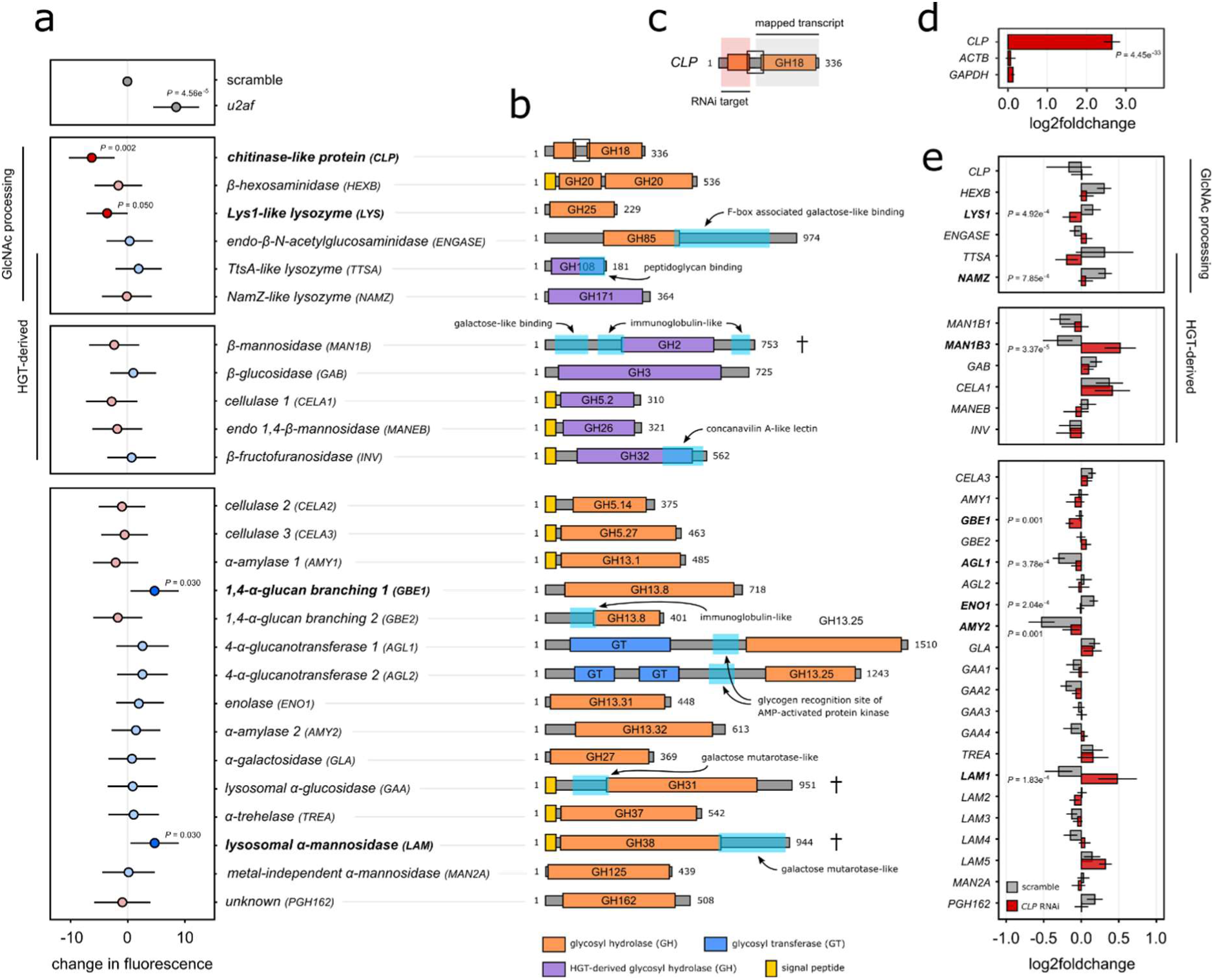
RNAi of a subset of glycan-processing genes perturbs host-mediated endosymbiont destruction, and perturbation of *CLP* determines expression of glycan-processing genes. All experiments were conducted using *Pb* cultures containing 75-100 cells (*a*) or 250,000 (*d, e*) cells. Cultures were fed daily with HT115 *E. coli* expressing target dsRNA to induce RNAi, or a non-hit ‘scramble’ dsRNA control. Per-host fluorescence is used throughout as a proxy for endosymbiotic algal load. *a*, Change in per host fluorescence, compared to scramble treatment, in RNAi treated *Pb* cultures (*n* = 12) during endosymbiont break-down. Cells were fed under standard conditions for 4 days, followed by 2 days of simultaneous RNAi feeding and cycloheximide treatment (50 µgmL^-1^) to trigger endosymbiont digestion. *b*, Schematic illustration of functional domain architecture of host putative glycosyl hydrolase (GH) encoding genes. For chimeric constructs (†) targeting multiple genes a single representative is shown (see **Fig. S3** for all genes). GH, glycosyl hydrolase (orange/purple); GT, glycosyl transferase (blue); additional domains are highlighted in cyan. *c*, Schematic illustration of *CLP* RNAi construct design including dsRNA target (red) and mapped transcript region for differential expression analysis (grey). *d*, Change in *CLP* gene expression (*c*) during RNAi treatment in untreated cultures, compared to Actin and GAPDH housekeeping genes. *e*, Change in GH transcript expression in scramble (grey) or *CLP* RNAi treated (red) *Pb* cultures (*n* = 6) during cycloheximide induced endosymbiont break-down, compared to respective scramble or *CLP* RNAi cultures without cycloheximide treatment (see also **Supplementary Table S2**). Statistical significance is shown for difference in log2foldchange values between scramble and *CLP* RNAi conditions. For all data, statistical significance was calculated using a linear mixed effects model (*a*) or an un-paired two-tailed t-test (*d*, *e*). Conditions where (*a*) *p* < 0.05 or (*e*) Bonferroni-corrected *p* < 0.0015 are shown.

RNAi of one GH and one GH family (chimeric RNAi-constructs were designed to target multiple GH paralogues belonging to the same family; **Fig. S3**) led to higher per-host fluorescence compared to scramble controls (control fluorescence = 44.6 relative fluorescence units [RFU]) (**Fig 2a-b**), suggesting that perturbation of these genes stalls endosymbiont destruction. These putative GHs were annotated as a *1,4-α-glucan branching enzyme* (*GBE1*) proposed to function in glycogen biosynthesis (fluorescence = 49.3 RFU, increased 10.6%; *t*=2.2, *df*=228, *p*=0.028) and a family of *lysosomal-α-mannosidases* (*LAM*) predicted to cleave mannose (fluorescence = 49.3 RFU, increased 10.6%; *t*=2.2, *df*=272, *p*=0.030). RNAi of two other GHs led to greater reduction in per-host fluorescence compared to scramble controls (**Fig. 2a-b**), suggesting that perturbation of these genes may increase endosymbiont destruction. These included a putative *chitinase-like protein* (*CLP*) predicted to bind GlcNAc oligosaccharides (fluorescence = 38.3 RFU, reduced 14.2%; *t*=-3.1, *df*=291, *p*=0.002) and a putative *Lys1-like lysozyme* (*LYS1*) predicted to cleave GlcNAc-GlcNAc (e.g., chitin) or GlcNAc-MurNAc (e.g., peptidoglycan) containing substrates (fluorescence = 41.0 RFU, reduced 8.1%; *t*=-1.97, *df*=290, *p*=0.050). The greater reduction in fluorescence observed upon *CLP* RNAi is intriguing because an orthologue of this gene binds to glycan moieties such as those present in *Chlorella* cell walls, and has been suggested to play a role in the sensing of chitin-containing microbes, amongst other functions^60–62^ (described in more detail below). We hypothesize that CLP may act as a binding protein which functions in a glycan processing/sensing signaling cascade involved in the regulation of endosymbiont load and therefore sought to further explore the role of *CLP* during endosymbiont destruction in *Pb*.

If *CLP* functions in a signaling pathway which orchestrates endosymbiont control, *CLP* RNAi perturbation should influence the transcription of other *Pb* encoded GH genes. To explore whether GH function was altered during endosymbiont destruction in a *CLP-*dependent manner, we assessed GH expression in cycloheximide treated and untreated control cultures, and compared changes in expression under *CLP* RNAi or scramble control RNAi conditions. Ensemble RNA sequencing was conducted under each condition and reads were mapped to the *Pb* 186b genome^19^. To confirm *CLP* perturbation in response to RNAi, reads were mapped only to the 3’ *CLP* transcript end to avoid cross-talk from the *CLP* RNAi construct (**Fig. 2c**). Importantly, *CLP* RNAi in untreated cultures resulted in 2.5-fold over-expression of *CLP* (**Fig. 2d**), suggesting either that compensatory gene expression was occurring in response to increased *CLP* dsRNA and subsequent siRNA exposure^16,63^, and/or that the RNAi signal (siRNA generation) had spread along the *CLP* transcript due to RNA-dependent RNA-polymerase amplification of cleaved mRNA transcripts that are over-represented in the mapped reads^16,64^. Both outcomes are known phenomena in *Paramecium* RNAi experiments^16,63,64^ and could interact, producing complexity in terms of phenotypic outcome. Regardless, *CLP* RNAi resulted in significant perturbation in *CLP* transcript abundance (**Fig. 2d**).

*CLP* RNAi during endosymbiont destruction significantly altered the expression of 8 (24%) GH genes (Bonferroni corrected since these did not test an *a priori* hypothesis; *p* < 0.0015) (**Fig. 2e**; **Fig. S4**). These include two putative GlcNAc-processing genes (*LYS1*; **Fig. 2a**, and *NAMZ*; **Fig. 1b**) and two putative HGT-derived genes (aforementioned *NAMZ*, and *MAN1B3*; **Fig. 1a**). The most significant changes in directionality of expression fold-change during endosymbiont destruction, resulting from *CLP* RNAi, were observed for *LYS1* (putatively involved in endosymbiont destruction; **Fig. 2a**), *MAN1B3* (a putative HGT-derived gene predicted to cleave mannose; **Fig. 1b**), and *LAM1* (within a GH family putatively involved in endosymbiont destruction; **Fig. 2a**). Interestingly, all three additional genes implicated in endosymbiont control upon RNAi (*LYS1, GBE1* and *LAM1;* **Fig. 2a**) were significantly altered upon *CLP* RNAi during endosymbiont destruction. These data demonstrate that a functional network of glycan-processing genes, at least in part controlled by *CLP*, acts in the regulation of endosymbiont destruction in *Pb*.

### CLP RNAi alters expression of transcription factors and intracellular signaling systems

To explore the broader role of *CLP* in *Pb*, we investigated how perturbed *CLP* function influenced additional pathways triggered during endosymbiont destruction. Untargeted metabolomics of *Pb* during *CLP* RNAi, compared to scramble control RNAi, revealed enrichment in metabolites belonging to cell signaling (inositol phosphate, and phosphatidylinositol signaling) and carbohydrate metabolic pathways (**Fig. 3a**; see also **Supplementary Table S3**). A significant reduction in relative abundance of several metabolites was detected upon *CLP* RNAi, including inositol 1,3,4-trisphosphate, fructose 1,6-bisphosphate, and unspecified glucan disaccharides, which may include maltose, the product of photosynthesis released by the algal endosymbionts^21–23^ (**Fig. 3b**; **Fig. S5**).

**Figure 3:**
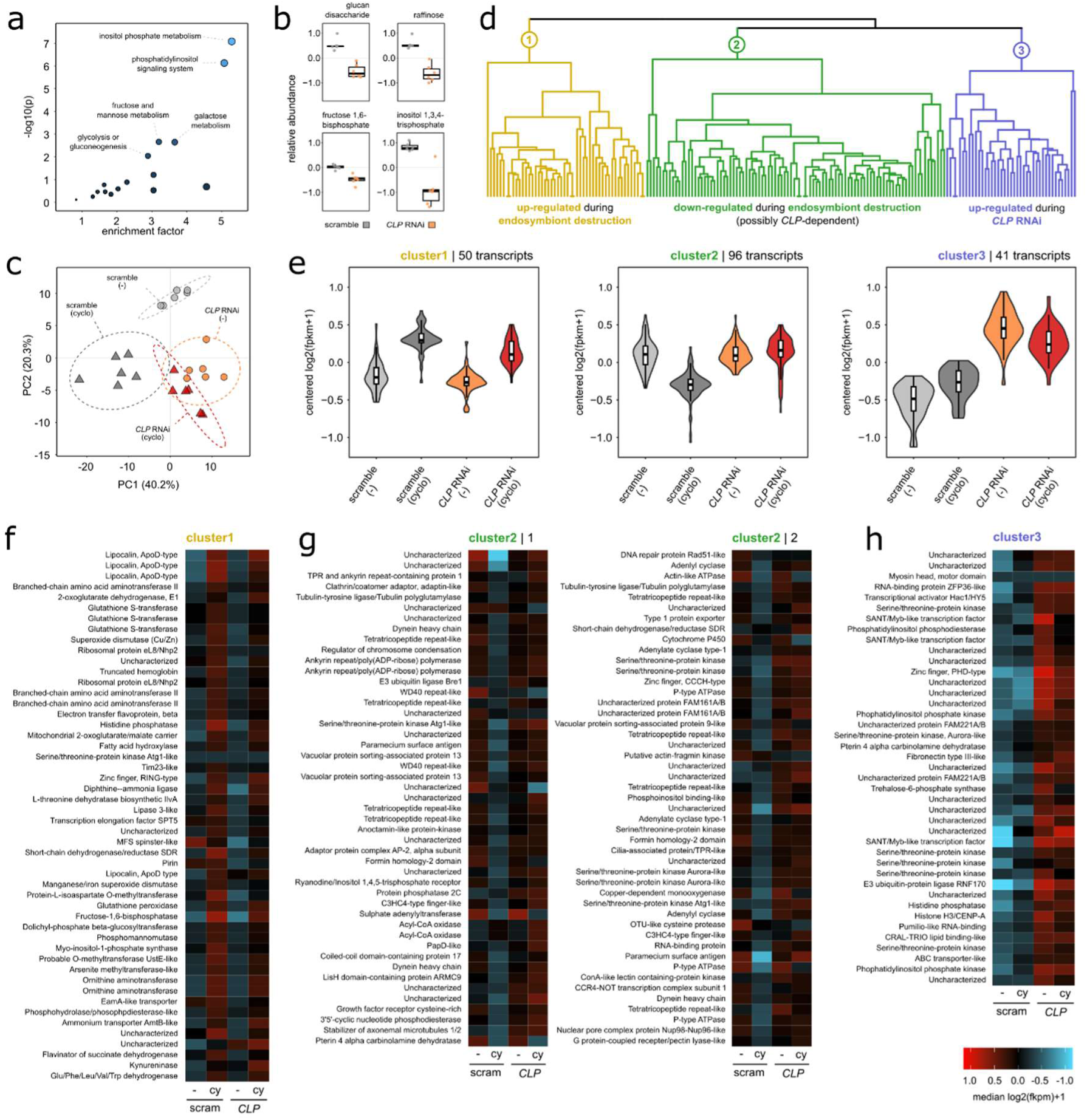
*CLP* RNAi alters expression of transcription factors and intracellular signalling systems. All experiments were conducted using *Pb* cultures (*n* = 6) containing approximately 250,000 cells. Cultures were fed daily with HT115 *E. coli* expressing target dsRNA to induce *CLP* knock-down, or ‘scramble’ dsRNA control with no hits to sampled *Pb* genes*. **a,*** Enrichment plot of significantly altered pathways during *CLP* RNAi, compared to a scramble control, detected by untargeted metabolomics (see also **Supplementary Table S3**). ***b,*** Relative abundance of significantly altered metabolites detected during *CLP* RNAi. ***c,*** PCA plot showing variance of transcriptional profiles in *Pb* cultures during *CLP* RNAi (orange, red) compared to a scramble control (grey, dark grey). Samples were treated with 50 µgmL^-1^ cycloheximide (triangle; cyclo) to trigger endosymbiont break-down, or remained untreated (circle; -). The percentage of explained variance for each PC is displayed. ***d,*** Hierarchical clustering of differentially expressed transcripts ([log2 fold change] ≥ 1, adjusted p value < 0.001) in each pairwise comparison. Clusters were defined through a dendogram maximum height cut-off of 80%. ***e,*** Three clusters of transcripts with similar expression patterns resulting from the cut dendogram (see also **Supplementary Table S4**) Data represented as median values (*n* = 6) of all transcripts from each cluster with overlaid distribution density. ***f-h,*** Heatmap of read-centred expression in each sample condition, and transcript annotation (InterProScan), for each of the identified transcript clusters. For relative abundance of all identified significantly altered metabolites during *CLP* RNAi, see **Fig. S4**. For individual heatmaps of each pairwise comparison across replicates during differential expression analysis, see **Fig. S6**.

We then assessed the expression of all *Pb* transcripts during endosymbiont destruction (cycloheximide treatment) and *CLP* RNAi, compared to both untreated and scramble control RNAi conditions (as per **Fig. 2e**). This identified distinct transcriptional profiles for each condition (**Fig. 3c**), with clear separation observed between untreated and cycloheximide treated cultures under scramble control RNAi conditions. A less distinct separation profile was observed between untreated and cycloheximide treated cultures during *CLP* RNAi, consistent with our hypothesis that functions associated with endosymbiont destruction may be coordinated by *CLP*. To explore transcriptional patterns associated with each treatment, we performed differential expression analysis for each pairwise condition. Differentially expressed transcripts ([log_2_ fold change] ≥ 1, adjusted p-value < 0.001) were hierarchically clustered based on relative expression under each condition (**Fig. 3d; Fig. S6**), yielding three distinct transcript clusters (**Fig. 3e**; see also **Supplementary Table S4**).

Cluster 1 contains transcripts that are up-regulated during cycloheximide treatment, independently of *CLP* RNAi (**Fig. 3f**), and includes genes related to autophagy (ser/thr-protein kinase Atg1), intracellular destruction and transport (lipocalin, lipase), carbohydrate metabolism (fructose-1,6-bisphosphatase; consistent with the mass spectrometry analysis, **Fig. 3b**), and oxidative stress (glutathione peroxidase). We propose that these are genes involved in the phago-lysosomal processes activated during destruction of algal endosymbionts^25–29^.

Cluster 2 contains transcripts predominantly down-regulated during cycloheximide treatment in scramble control cultures (**Fig. 3g**), and so represent host functions possibly inactivated during endosymbiont destruction. Interestingly, these transcriptional patterns were not observed upon cycloheximide treatment coupled with *CLP* RNAi (**Fig. S6c-d**), supporting the proposition that *CLP* RNAi can perturb endosymbiont control (seen in **Fig. 2a**). This cluster includes genes predicted to function in signal transduction, and two putative phosphatidylinositol binding proteins (again consistent with the metabolomics data; **Fig. 3a**) indicating that phosphatidylinositol signaling is altered during regulation of endosymbiont destruction. Phosphoinositides – a broad class of membrane phospholipids – are key regulators of endosomal processes and downstream signals of extracellular receptor activation, acting in intracellular membrane trafficking, autophagy, and innate immunity^65–69^. Other genes in this cluster were predicted to be involved in intracellular vesicular trafficking (vacuolar sorting-associated proteins 9 & 12, and adaptin-like proteins) and cytoskeletal arrangement (dynein heavy chain, and tubulin-tyrosine ligase/polyglutamylase). We propose that these genes function in the cellular re-arrangements that maintain endosymbiosis in *Pb*, and may represent processes that are inactivated during endosymbiont destruction, such as detachment of the perialgal vacuole from the host cell cortex and lysosomal fusion^25–29^. The observed down-regulation of these genes during algal destruction is consistent with the proposal that these systems function in maintenance of the endosymbiotic population.

Cluster 3 contains transcripts with significantly increased expression during *CLP* perturbation (**Fig. 3h**), irrespective of cycloheximide-induced endosymbiont destruction (as in Clusters 1 and 2). This includes genes predicted to function in signal transduction, including phosphatidylinositol signaling (as in **Fig. 3g**), serine-threonine protein kinases, transcription factors, and a transcriptional activator. We therefore propose that these may be components of a signaling cascade in which *CLP* is a component, and suggest that disruption of this signal through *CLP* RNAi may be responsible for the increased endosymbiont destruction trend observed during cycloheximide treatment (**Fig. 2a**). Taken together, these findings indicate that complex signaling mechanisms are involved in host-mediated endosymbiont destruction, and suggest that *CLP* may act as a key regulator of these molecular processes.

### CLP localizes proximate to the putative host phagosome and lysosome during endosymbiont destruction

To understand the role of *CLP* in endosymbiont regulation, we performed immunofluorescence microscopy to track CLP localization during cycloheximide-induced endosymbiont destruction. Using a custom polyclonal antibody raised against *Pb* CLP (see **Methods**), we observed CLP localization to the host phagosome and oral apparatus (**Fig. 4a**; see also **Supplementary Table S5**). These cell systems are part of the ciliate feeding grove^70,71^ and are associated with both heterotrophic feeding and establishment/regulation of endosymbiotic algae^17,18^. A large fraction of the cells exhibited phagosome-like CLP localization (78.4% of cells in untreated conditions, 85.7% in cycloheximide treated). However, some cells demonstrated an additional pattern of CLP localization consistent with host lysosomes (32.4% of cells in untreated conditions, 40.3% in cycloheximide treated; **Fig. 4b**). These observations indicate that *CLP* is involved in at least two systems (phagosomal and lysosomal) which have complex overlapping functions. These data also demonstrate that the sampled population includes cells which are at mixed stages of response, as identified by differential CLP localization patterns. This result is consistent with host cell and associated endosymbiotic functions not being synchronized (e.g., only 32.4% of cells demonstrating CLP-lysosome localization and 78.4% of cells demonstrating CLP-phagosome localization under standard conditions; **Supplementary Table S5**).

**Figure 4:**
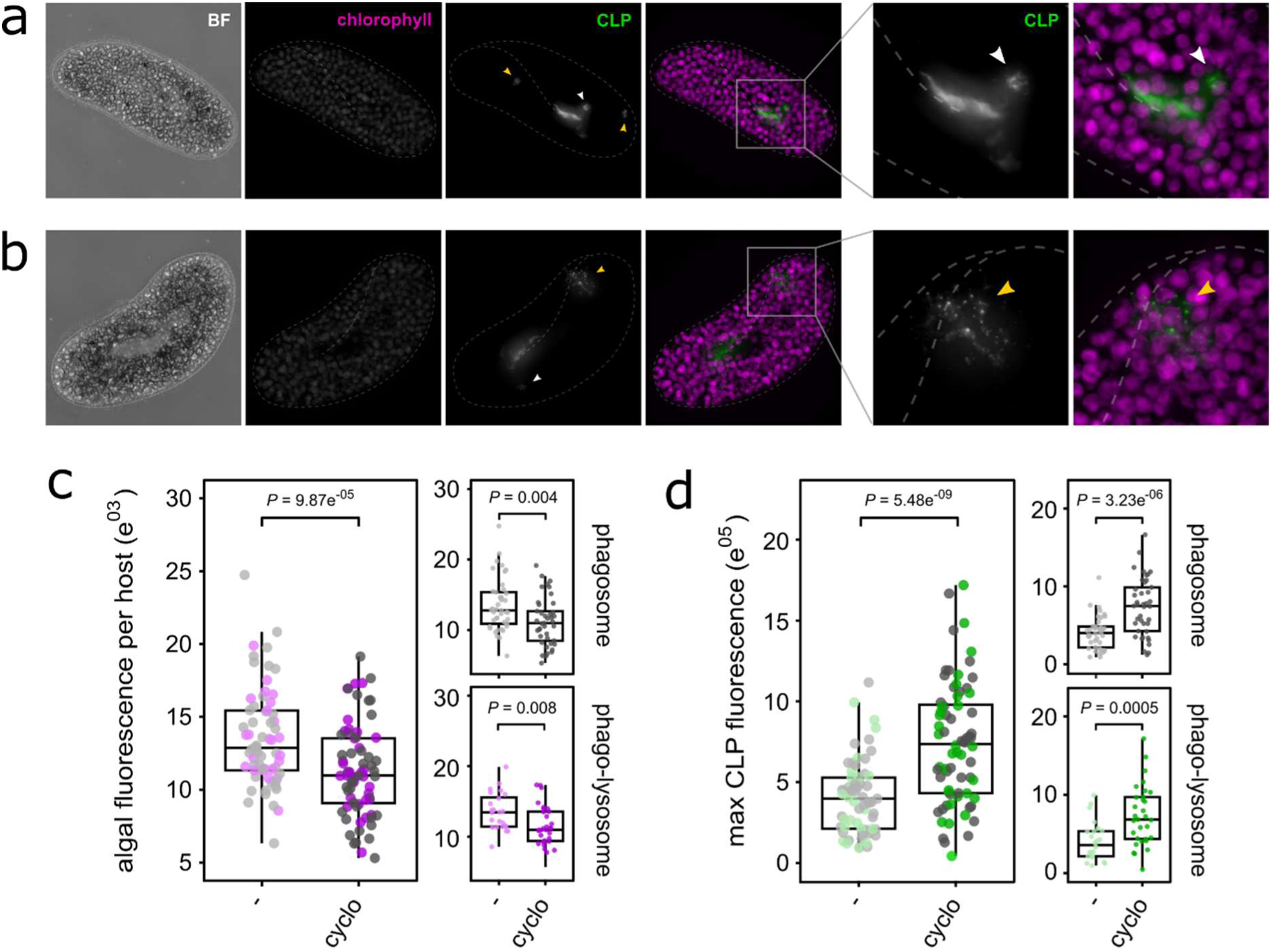
CLP localizes proximate to the putative host phagosome and lysosome during endosymbiont destruction. All experiments were conducted using *Pb* cells grown under standard conditions (- ; *n* = 70), or treated with 50 µgmL^-1^ cycloheximide (cyclo ; *n* = 69). All cells were treated a primary αCLP antibody followed by an Alexa Fluor 488-conjugated secondary antibody for immunofluorescence imaging. ***a-b***, Fluorescence imaging of *Pb* cells exhibiting phagosome-like (***a*** ; white arrows) or lysosome-like (***b*** ; yellow arrows) CLP localization. Magenta indicates endosymbiotic algal autofluorescence, green indicates Alexa Fluor 488 secondary antibody fluorescence. ***c-d*** Mean algal fluorescence (***c***) and max CLP fluorescence (***d***) per host cell in untreated (-) or cycloheximide treated (cyclo) cells. Cells were scored based on whether they exhibited phagosome- or phagosome/lysosome-like *CLP* localization. Boxplot data are represented as max, upper quartile (Q3), mean, lower quartile (Q1) and min values, and individual data points for each biological replicate are shown. Statistical significance was calculated using a linear model.

To explore how CLP localization is adjusted during endosymbiont destruction, we compared algal autofluorescence and CLP immunofluorescence in *Pb* during cycloheximide-induced endosymbiont destruction. A significant reduction in per-host algal fluorescence during cycloheximide treatment (**Fig. 4c**) was coupled with increased CLP fluorescence (**Fig. 4d**). This trend was observed for both phagosome and phago-lysosome-like exhibiting cells, indicating that endosymbiont destruction is coupled to *CLP* dynamics in both cellular processes. These data support the hypothesis that *CLP* is functioning as a sensor for the detection of endosymbiont or food-derived (**Fig. 4a-b**) glycan-substrates released during endosymbiont destruction and heterotrophic feeding, and is consistent with data suggesting the *LYS1* lysozyme (predicted to liberate these substrates and which is regulated by *CLP*) is involved in endosymbiont destruction (**Fig. 3a, e**).

### CLP is a chitin-binding protein homologous to an animal immune factor

To further explore the function of *CLP* in *Pb,* we sought to understand the evolution of this gene family across the eukaryotes. A single homologue of *Pb CLP*, *Stabilin-interacting Chitinase-like Protein* (*SI-CLP*), was identified in the human genome. *SI-CLP* is an innate immune-factor expressed in macrophages and neutrophils that activates signal-cascades involved in inflammatory responses^62^ (including phosphoinositide kinases^72^; **Fig. 3a, f-h**), and is sorted into CD63+/LAMP1-positive lysosomes and p62lck-positive late endosomes in mammals, indicating a role in secretory, digestive and autophagy-related intracellular trafficking^61^. *SI-CLP* has broad substrate-binding capacity that includes GlcNAc oligosaccharides and other glycan moieties, and may play a sensory role in host responses to chitin-containing pathogens^60^. No homologue for the *stabilin-1* protein, with which human *SI-CLP* interacts, was identified in *Pb* or other ciliates (NCBI protein database; gathering threshold: e-01; June 2024) and so this protein was not explored further.

Phylogenetic reconstruction of 1,445 candidate homologues reveals that *CLP* is present in the majority of extant eukaryotic phyla searched (**Supplementary Dataset S2**) demonstrating that this is an anciently-derived eukaryotic gene family. An amino acid alignment of representative eukaryotic *CLP* homologues (**Fig. 5a**; **Fig. S7**) revealed strong conservation of binding sites (all but one retained in 90% of the homologues aligned) and predicts the presence of a 5’ secretion signal peptide sequence in 75% of the sequences sampled (DeepLoc2.0)^73^. Identifiable chitin cleavage sites^60,61^ were not conserved, with universal absence of the 2^nd^ and 4^th^ known catalytic residues, and 78% of species compared lacking all but one known catalytic residue. Putative binding and cleavage sites were mapped to the predicted structure of *Pb* CLP (AlphaFold3; May 2024) revealing a pore-like structure in which GlcNAc oligosaccharide binding is proposed to occur (**Fig. 5b**). Comparison with the crystal structure of *H. sapiens (Hs)* SI-CLP^60^ (which forms a dimer; **Fig. S8**) reveals positional similarity of binding and cleavage sites. Western blot analysis of *Pb* CLP reveals two bands close to 37 kDa and 75 kDa (predicted molecular weight of *Pb* CLP, 40.24 kDa; see **Fig. S9**) indicating that the *Pb* homologue may also form a dimer.

**Figure 5:**
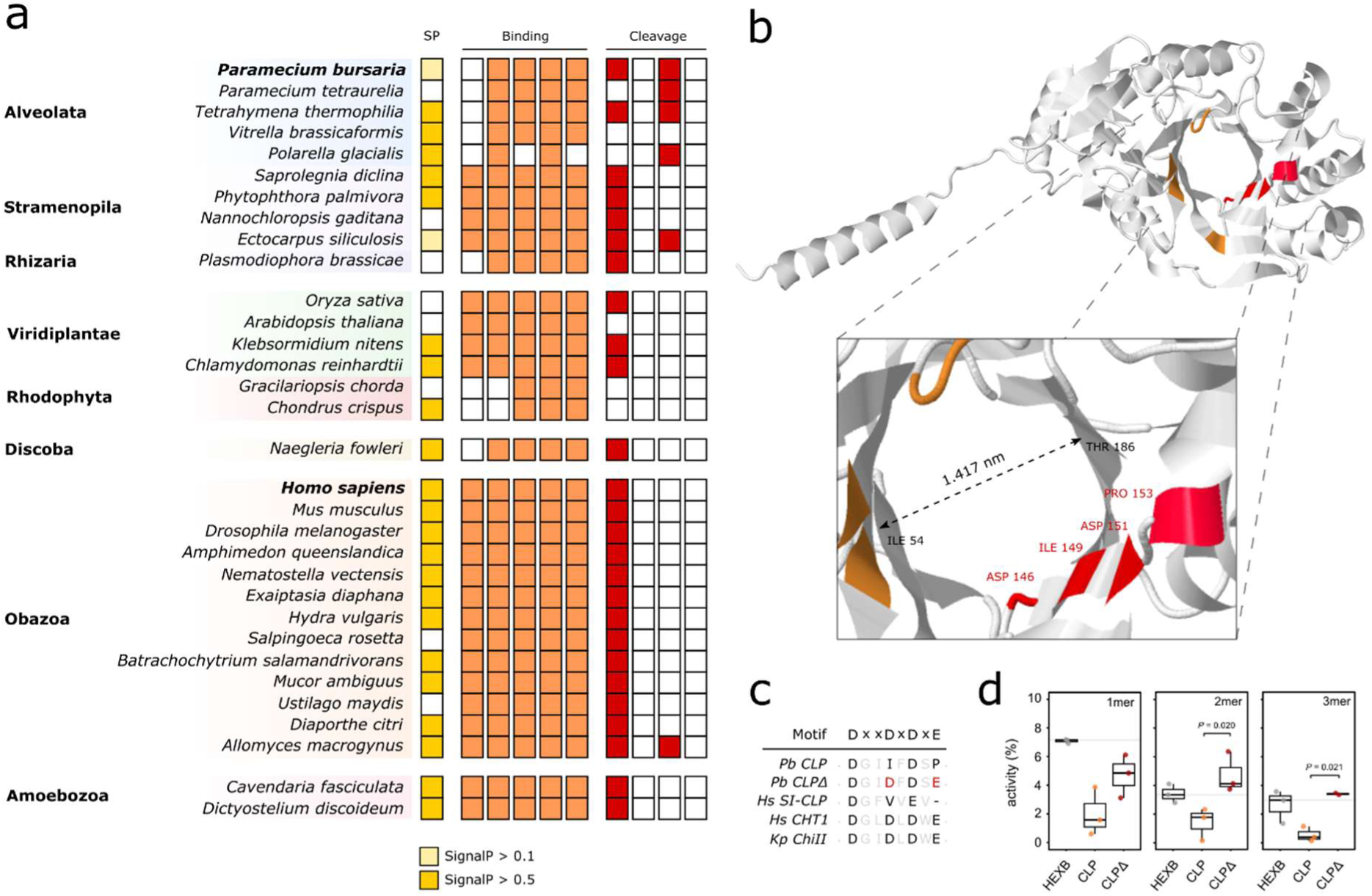
CLP is a chitin-binding protein homologous to an animal immune factor. **a,** Conserved functional domain residues of *CLP* across a sample of eukaryotic protein homologues (see also **Fig. S7**) including predicted signal peptide (yellow) and characterised binding (orange) and cleavage (red) site amino acids. ***b,*** Predicted 3D structure of *Pb* CLP generated with AlphaFold3 (see also **Fig. S8**). Characterised binding (orange) and cleavage sites (red) are highlighted. ***c,*** Trimmed amino acid alignment of *Pb CLP* and *Hs SI-CLP* sequences, compared to known chitin-cleaving enzymes *Hs CHT1* and *K. pneumoniae ChiII*. Residues in bold are sites associated with a conserved chitin-cleavage motif. Residues in red indicate synthetically mutated sites to ‘restore’ chitin-cleaving function. ***d.*** Percentage activity of purified wild-type (CLP) and mutated (CLPΔ) *Pb* enzymes incubated for 3 hours in 1mer (GlcNAc), 2mer (GlcNAc_2_; Chitin_2_), or 3mer (GlcNAc_3_; Chitin_3_) chitin substrates. Enzymes were synthesized to encode a 5’ yeast secretion signal, transformed into chitinase-deficient *Saccharomyces cerevisiae* strain Y16947, and expressed under selection for 7 days. Supernatant containing the expressed enzyme was concentrated, washed, and protein concentrations matched prior to the assay. Activity was standardised against supernatant from a yeast empty vector control. *Pb β-hexosaminidase* (Fig. 2b), an enzyme predicted to cleave terminal GlcNAc residues from glycan chains, was used as a positive control (HEXB). Boxplot data are represented as max, upper quartile (Q3), mean, lower quartile (Q1) and min values, and individual data points for each biological replicate are shown. Statistical significance was calculated using generalized linear models with quasi-binomial error distribution.

*Pb CLP* and *Hs SI-CLP* lack the conserved ‘DxxDxDxE’ motif associated with chitin-cleavage (**Fig. 5c**). To understand catalytic function, *Pb* CLP engineered to carry a *Saccharomyces* secretion peptide was expressed in a chitinase-deficient *Saccharomyces cerevisiae* strain, and the secretome assayed using a Chitinase Activity Kit (**Fig. 5d**). We observed low catalytic activity of *Pb* CLP on all substrates compared to *Pb* HEXB (predicted to cleave terminal GlcNAc residues from glycan chains; **Fig. 2b**). It is important to note that low activity of all enzymes, including HEXB, may be due to the required assay conditions (37°C) being sub-optimal for *Pb* enzymes (∼18-25°C). Nonetheless, we observed that directed mutation of the 2^nd^ and 4^th^ active site residues of *Pb CLP* (*CLPΔ;* **Fig. 5c**) ‘restored’ cleavage function and significantly increased activity on 2mer (GlcNAc_2_ / Chitin_2_) (*t*=3.141; *df*=6; *p*=0.020) and 3mer (GlcNAc_3_ / Chitin_3_) (*t*=3.324; *df*=6; *p*=0.021) substrates (**Fig. 5d**), supporting the hypothesis that these residues are important for catalytic activity. These data indicate that *CLP* is a eukaryotic wide gene family which functions in GlcNAc oligosaccharide (e.g., chitin) binding, but lacks potent cleavage activity, supporting our hypothesis that *CLP* plays a role in glycan sensing.

### Glycan break-down products alter host control of endosymbiotic algae

Our genomic and RNAi data point to glycan-sensing/processing being central to host-mediated endosymbiont control. As such, we next explored how glycan break-down substrates were altered during regulation of endosymbiosis in *Pb*, and the reciprocal effect of these substrates on endosymbiont control. Here, we used altered light intensity to trigger endosymbiont destruction^13,30^ to simulate conditions experienced by *Pb* in nature. We observed a reduction in *Pb* per-host fluorescence upon exposure to high-light (80 μ mol m^-2^ s^-1^; **Fig. 6a**). This was accompanied by no comparable reduction in host cell number (**Fig. 6b**) indicating that these conditions did not have a negative effect on host viability during the period of the experiment. Analysis of metabolite abundance in *Pb* under varying light conditions (50 μ mol m^-2^ s^-1^ to 6 μ mol m^-2^ s^-1^)^30^ revealed that reduction in endosymbiont load was accompanied by increased abundance of GlcNAc and glucosamine (GlcN – a substrate which also binds to *SI-CLP*^60^) (**Fig. 6c**; **Fig. S10**). These substrates are consistent with break-down products of chitin-like glycans^32–34^ (**Fig. 6d**) and support the hypothesis that host control of endosymbiotic *Chlorella*^13,30^ is facilitated through break-down of glycans present in the algal cell wall. This is also consistent with the predicted function and involvement of *LYS1* (**Fig. 2a/e**) and the HGT-derived *NAMZ* (**Fig. 2e**) in endosymbiont destruction under *CLP*-derived transcriptional control.

**Figure 6:**
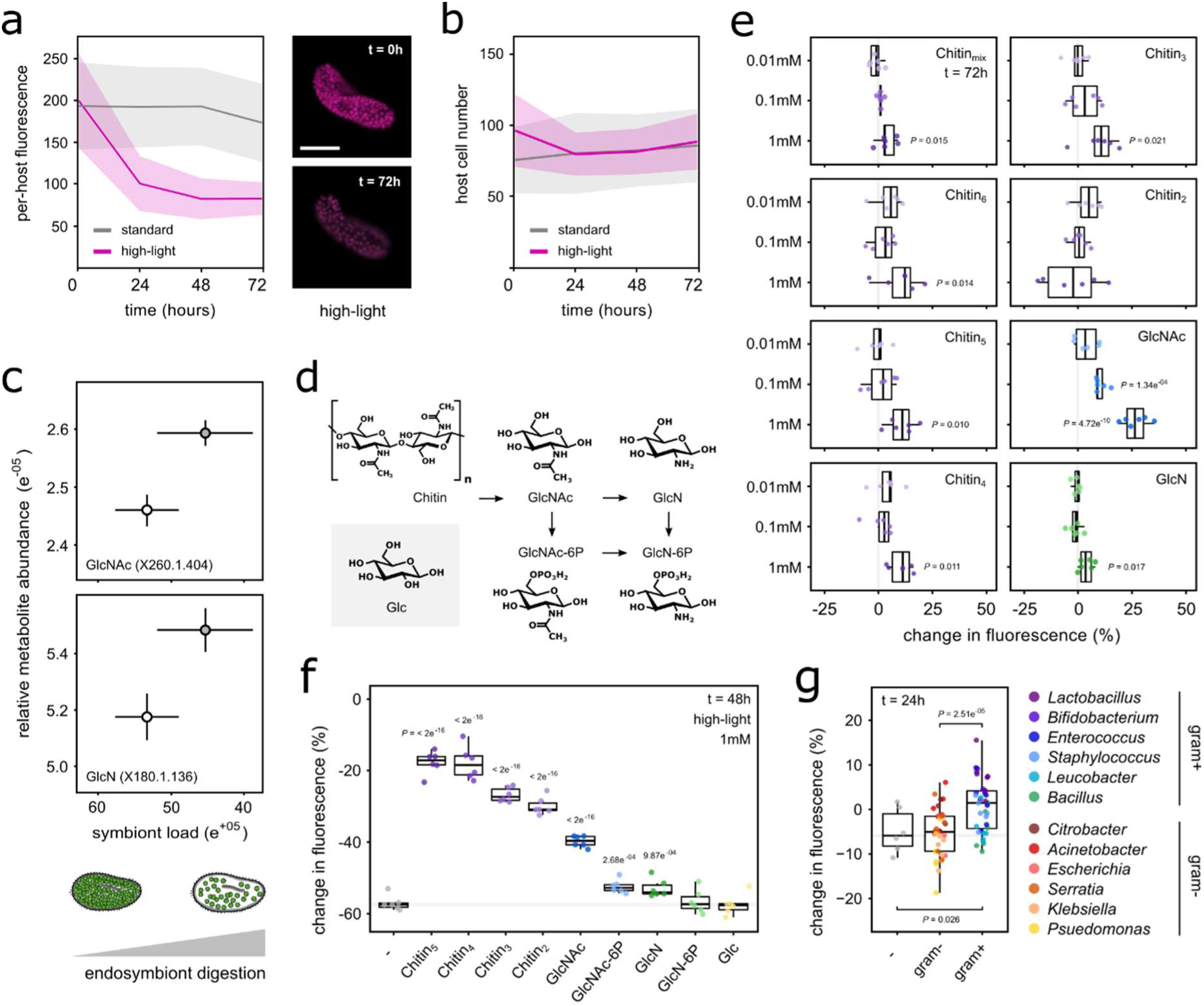
Glycan break-down products alter host control of endosymbiotic algae. All experiments were conducted using pre-starved *Pb* cultures (*n* = 6) grown under standard conditions (see **Methods**), unless stated otherwise. Growth under standard light was conducted at 25 μ mol m^-2^ s^-1^, high-light at 80 μ mol m^-2^ s^-1^, moderate high-light at 50 μ mol m^-2^ s^-1^, and low-light at 6 μ mol m^-2^ s^-1^. Per host fluorescence is used throughout as a proxy for endosymbiotic algal load. ***a***, Mean per host fluorescence in *Pb* cultures grown under high-light. Data are presented as mean ± sd. Scale bar = 50μm, magenta indicates endosymbiotic algal autofluorescence. ***b***, Host cell number in *Pb* cultures grown under high-light for 72 hours. Data are presented as mean ± sd. ***c***, Relative metabolite abundance in *Pb* cultures under variant light conditions, in which differential endosymbiont digestion is known to occur. Cells were grown under moderate high-light (white) or low-light (grey) for 3 days prior to metabolite extraction, and symbiont load per host-cell assessed by flow cytometry (**Fig. S10**). ***d***, Schematic metabolic pathway of chitin break-down, including the structure of glucose (Glc) for comparison. ***e***, Percentage change in per host fluorescence following exposure to chitin and chitin break-down products, compared to an untreated control. Cells were exposed to chitin (mixed oligosaccharides, or oligosaccharide lengths 6 to 2), GlcNAc, or GlcN for 72 hours. ***f***, Percentage change in per host fluorescence upon exposure to chitin break-down products during high-light initiated endosymbiont digestion. Cells were grown under high-light and exposed to chitin (oligosaccharide length 5 to 2), chitin break-down products (see ***d***), or Glc for 48 hours. Each substrate was added to a final concentration of 1mM. ***g***, Pooled percentage change in per host fluorescence following bacterial feeding in standard growth conditions. *Pb* cells were grown for 24 hours, and bacteria were heat-killed and matched to OD 0.1 prior to feeding (**Fig. S12**). All boxplot data are represented as max, upper quartile (Q3), mean, lower quartile (Q1) and min values, and individual data points for each biological replicate are shown. Statistical significance was calculated using a linear model on log-transformed data (**e**, **f**, **g**). Conditions where *p* < 0.05 are shown. GlcNAc, N-acetyl-D-glucosamine; GlcN, D-glucosamine; GlcNAc-6P, N-acetyl-D-glucosamine-6-phosphate; GlcN-6P, D-glucosamine-6-phosphate; Glc, Glucose.

Given the putative role of *CLP* as a glycan-sensor, we next tested how increased glycan abundance influenced endosymbiont control in *Pb*. We observed increased per-host fluorescence upon exposure to chitin, GlcNAc, and GlcN under standard growth conditions (**Fig. 6e**). This effect was greatest upon exposure to GlcNAc and remained significant at lower doses (0.1 mM; *t*=4.314; *df*=20; *p*=1.34^-04^), indicating sensitivity to this substrate which determined symbiotic outcomes. Supplementing separately cultured endosymbiosis-derived algae with GlcNAc did not enable heterotrophic growth or improve autotrophic growth compared to nitrogen free conditions (**Fig. S11a/b**), demonstrating that the response to chitin, GlcNAc, and GlcN seen in host was not due primarily to an algal growth response. Host exposure to chitin oligomers (*t*=18.860-24.805; *df*=50; *p* < 2e^-6^), GlcNAc (*t*=12.896; *df*=50; *p* < 2e^-6^), N-acetyl-d-glucosamine-6-phosphate (GlcNAc-6P) (*t*=3.922; *df*=50; *p*=0.004), and GlcN (*t*=3.5; *df*=50; *p*=0.010) during high-light treatment all significantly arrested the reduction in per-host fluorescence associated with endosymbiont destruction (**Fig. 6f**). Exposures were conducted at equal molar concentrations indicating that the length of the chitin polymer chain may be an important factor. No response was observed for D-glucosamine-6-phosphate (GlcN-6P) or glucose (Glc), indicating that this effect was not due to increased nutrient provision (carbon, nitrogen, or phosphate) to the host. Importantly, these GlcNAc mono- and oligosaccharides are consistent with substrates bound by *CLP* (**Fig. 5d**), and putative products of GlcNAc-processing GHs (e.g., *LYS1*) implicated in *CLP*-dependent endosymbiont destruction (**Fig. 2a/e**).

Finally, we sought to explore how broader mixotrophic functions could influence endosymbiont control. *Pb* is a mixotroph and phagocytoses bacteria in nature, therefore the host must be exposed to additional sources of glycans. We observed that exposure to heat-killed gram-positive bacteria, possessing exposed peptidoglycan-rich cell walls, also increased algal fluorescence per-host (*t*=4.492; *df*=75 ; *p*=2.51e^-05^) compared to heat-killed gram-negative bacteria delivered at the same dose (**Fig. 6g**; **Fig. S12**). This suggests that broader microbial trophic interactions, including mixotrophic feeding, may also influence the dynamics of endosymbiosis in *Pb*.

These data have revealed that exposure to GlcNAc, and other break-down products of glycans present in endosymbiotic algae and the cell walls of bacterial food, can alter endosymbiont control in *Pb.* We propose that *CLP*, which functions to bind these substrates, may act to regulate the broader cellular network that underpins this process. We therefore postulate that glycan-sensing and processing represents an important mechanism of endosymbiont control in *Pb* through the regulation of endosymbiont destruction.

## Discussion

Here we show that an immune-like glycan-sensing/processing network involving a conserved eukaryotic chitinase-like protein (*CLP*), and augmented by genes acquired through HGT, orchestrates the regulation of endosymbiosis in *Pb*. We demonstrate that host-mediated destruction of endosymbiotic algae releases glycan-substrates (**Fig. 6c**), specifically GlcNAc and GlcN, and that environmental exposure to GlcNAc mono- and oligosaccharides stalls endosymbiont destruction (**Fig. 6f**). Through comparative phylogenetics we reveal that *CLP* is related to an innate immune-factor in humans (*SI-CLP*) involved in phago-lysosomal response pathways and possible molecular-sensing of pathogens^60–62^, and show that *CLP* in *Pb* serves a similar function to coordinate phago-lysosomal activity. *CLP* localizes to the oral apparatus, phagosome, and lysosome of *Pb* with increasing localization intensity during endosymbiont destruction (**Fig. 4d**), where GlcNAc-substrates arise as they are liberated through endosymbiont destruction (**Fig. 6c**) or bacterial feeding (**Fig. 6g**). We propose that these substrates are acted upon by at least six ancient eukaryotic genes (e.g., *LYS1* - a GH that directly determines the fate of algal endosymbionts; **Fig. 2a**) and two horizontally-acquired glycan-processing genes (e.g., *MAN1B1, NAMZ;* **Fig. 1b**) which are transcriptionally regulated by *CLP* function (**Fig. 2e**). We hypothesize that this mechanism of glycan-sensing through *CLP* triggers a network of cellular processes involving transcriptional regulation, intracellular signaling (**Fig. 3f**-**h**) and glycan-processing (**Fig. 2e**) to regulate endosymbiont destruction in *Pb*.

Intriguingly, this host network of cellular processes includes genes acquired by *Pb* through prokaryote-to-eukaryote HGT. Among the genes altered through *CLP* perturbation during endosymbiont destruction are two genes proposed to be HGT-derived. *MAN1B3* (**Fig. 2a**), a predicted β-mannosidase, may be important for destruction of mannose-substrates in the endosymbiont cell wall. However, *NAMZ* (**Fig. 2b**), a predicted lysozyme, is potentially the most relevant, as this enzyme is proposed to cleave GlcNAc in chitin (i.e., from the endosymbionts cell wall) or peptidoglycan (i.e., from the bacterial cell wall) substrates. The interaction of predicted cleavage products of HGT-derived *NAMZ* (i.e., GlcNAc) with *CLP* (**Fig. 5d**) and their effect on endosymbiont destruction dynamics in the host (**Fig. 6e, f**), combined with the finding that *CLP* perturbation influences subsequent expression of *NAMZ* (**Fig. 2e**), suggests that this may represent part of the feedback loop of glycan-processing and sensing proposed to occur during endosymbiont destruction. These observations demonstrate how recently acquired gene functions (e.g., *NAMZ*) following HGT can augment ancestral functions (*CLP*).

‘You are what you eat’ is a hypothesis which posits that phagocytosis can drive HGT, from consumed cells to host nuclei, through establishment of a gene-transfer ratchet^74^. This process has been posited as an important factor in endosymbiotic organelle acquisition during eukaryotic evolution. The ‘shopping bag’ model went further, stating that the step-wise acquisition of genes from former, phagocytosed cells (i.e., food), sometimes involving cells which formed transient endosymbioses, built the host genetic repertoire required for endosymbiotic functions that, in turn, allows for stable symbiont integration^75,76^. This process is thought to be key for the evolution of plastid (chloroplast) organelles (i.e., eukaryotic photosymbiosis). In *Pb*, we observe data consistent with both hypotheses relating to the function of a facultative endosymbiosis, but with a fundamental addition. Here, the genes acquired by *Pb* arose not from transient interactions with eukaryotic algae, but from bacteria. The occurrence of bacterial genes in eukaryotic genomes is not uncommon^54^, though few cases have been reported in ciliates^77–80^ but also involve the transfer of GH genes, supporting the idea that genes that function in glycan-processing are prime candidates for prokaryote-to-ciliate HGT. *Paramecium* are well known for their ability to phagocytose or form stable symbiotic interactions with a diverse consortium of bacterial species^81–84^. Thus, we hypothesize that these genes likely originated from ancestral bacteria maintained either stably or transiently in endosymbiosis^85,86^, or ingested as food^74,76^. Regardless, our discovery that these genes of bacterial origin may be involved in endosymbiotic algal destruction in *Pb* illuminates how cross-kingdom HGT can shape host-microbe endosymbiotic interactions consistent with both the ‘you are what you eat’ and the ‘shopping bag’ hypotheses.

The role of innate immunity in plant-microbe and animal-microbe symbioses is well-understood^44,46,47,87^, however such mechanisms have yet to be described in a single-celled endosymbiotic system^88,89^. *Pb* lacks many components of a ‘canonical’ immune system (**Supplementary Table S6**). Yet detection of self from non-self must be critical to allow the host to defend against harmful microbial pathogens^90–92^, and mechanisms of molecular sensing that can determine friend from foe, or ‘good’ symbionts from ‘bad’, are essential for a host maintaining a population of intracellular symbionts^5,7,12–15,87,93^. This is important when the fitness interests of symbiont and host are not aligned, resulting in conflict between partners – an inevitability in symbiosis^94^– and whereby detection and resolution of conflict are needed for long-term stability of an interaction^13,29,31,95^. In *Pb*, maintenance of a population of endosymbiotic algae requires significant re-modelling and co-opting of host cellular machinery^18,25–27^. As such, a finely tuned network of molecular processes is required that allows the host to detect context-dependent cues, and respond to, initiate, maintain, or terminate the interaction. Here, we propose that this process occurs through the sensing of glycan-substrates released during endosymbiont destruction or microbial feeding (**Fig. 6a-f**), detected by *CLP*, and subsequent signaling that prompts the host to stall further endosymbiont destruction. This process would act to minimize the costly effects of mass endosymbiont destruction^29^, perpetuating surviving endosymbionts, and promoting long-term stability of the interaction. Our discovery highlights how a putatively ancient mechanism of immune-like glycan sensing, shaped partly by functions acquired via prokaryote-to-eukaryote HGT, can be co-opted for control of a facultative endosymbiotic interaction. We propose that this functional network of glycan processing and sensing represents an important factor in how stable host-microbe associations have evolved.

## Data Availability

All data and script used for analysis and processing are available at: https://github.com/benjaminhjenkins/Paramecium_bursaria. All sequence reads are available on NCBI GenBank with the BioProject identifier PRJNA65904 and SRA accession number SRR1251101, and at: https://github.com/guyleonard/paramecium and https://zenodo.org/doi/10.5281/zenodo.4638887.

## Acknowledgements

This work was primarily supported by an ERC Consolidator Grant (CELL-in-CELL), Gordon and Betty Moore Foundation (GBMF) grant (GMBF11490) and a Royal Society University Research Fellowship (UF130382) to T.A.R.. MAB and DDC were supported by grants from the BBSRC (BB/X016439/1) and NERC (NE/V000128/1). We thank Prof. Neil Gow of the University of Exeter for informative discussions on chitin sensing which inspired this work. The authors gratefully acknowledge the Micron Advanced Bioimaging Facility (supported by Wellcome Strategic Awards 091911/B/10/Z and 107457/Z/15/Z) for their support & assistance in this work.

## Methods

### Culture conditions and media

In all experiments, *Paramecium bursaria* (*Pb*) 186b (CCAP 1660/18) strain was used. *Pb* cells were cultured in New Cereal Leaf – Prescott Liquid media (NCL), unless modified as described. NCL media was prepared by adding 4.3 mgL^-1^ CaCl_2_.2H_2_O, 1.6 mgL^-1^ KCl, 5.1 mgL^-1^ K_2_HPO_4_, 2.8 mgL^-1^ MgSO_4_.7H_2_O to deionised water. 1 gL^-1^ wheat bran was added, and the solution boiled for 5 minutes. Once cooled, media was filtered once through Whatman Grade 1 filter paper and then through Whatman GF/C glass microfiber filter paper. Filtered NCL media was autoclaved at 121°C for 30 mins to sterilise prior to use.

NCL medium was bacterized with *Klebsiella pneumoniae* SMC and supplemented with 0.8 mgL^-1^ β-sitosterol prior to propagation. *Pb* 186b cells were sub-cultured 1:9 into fresh bacterized NCL media every two months and maintained at 23°C at a light intensity of 25 = μ mol m^-2^ s^-1^ (standard – e.g., **Fig. 6a/b**) or 80 μ mol m^-2^ s^-1^ (high – e.g., **Fig. 6a/b**) with a light-dark (LD) cycle of 12:12 h. *Pb* cultures were maintained for at least three weeks following sub-culture to ensure that cells were starved and in stationary phase prior to experimentation.

In free-living algal assays, *Micractinium conductrix* SAG 241.80 (a member of the *Chlorella* clade^20,22^) was used, as axenic algal endosymbionts isolated from *Pb* 186b could not be reliably cultured. This strain was isolated from *Pb* SAG 27.96 and shares an identical ITS2 sequence with the *M. conductrix* endosymbiont of *Pb* 186b^96^. Algae were cultured in Modified Bold’s Basal Medium (MBBM) medium at 24°C with a 14:10 h LD cycle.

For bacterial feeding assays the following strains were used: *Escherichia coli* HT115 (Ec); *Klebsiella pneumoniae* SMC (Kp); *Serratia marascens* BS303 (Sm); *Leucobacter celer subsp. astrifaciens* CBX151T (Ls); *Actinetobacter baumanii* (Ab); *Bacillus subtilis* 168 (Bs); *Psuedomonas aeruginosa* PA14 (Pa); *Enterococcus faecalis* OG1RF (Ef); *Staphylococcus aureus* MSSA476 (Sa); *Citrobacter freundii* (Cf); *Bifidobacterium breve* (Bb); and *Lactobacillus ruminis* (Lr). Bacteria were cultured overnight (constant shaking, 30℃) in Luria-Bertani broth (Ec – Pa) or Todd-Hewitt (Ef – Sa) broth based on standard cultivation methods, or in modified Gifu Anaerobic Medium (mGAM; Nissui Pharmaceuticals) (Cf – Lr) under anaerobic conditions^97^ (5% H2, 5% CO2, 90% N2, <20 ppm O2).

For yeast secretion assays, *Saccharomyces cerevisiae* Y16947 (Euroscarf) was used. This strain is a *S. cerevisiae* BY4742 CTSI deletion mutant (YLR286cΔ). *S. cerevisiae* was cultured in YPD at 30°C and stored at -80°C in 25% glycerol solution.

### Phylogenetic analysis

Dataset generation and sampling for each phylogenetic analysis were conducted as follows. For identification of CAZymes in *Pb*, predicted proteins^19^ were searched with a custom annotation tool (https://github.com/benjaminhjenkins/CAZyme_survey). For HGT analysis, dataset curation was conducted as described by Irwin et al.^98^. Briefly, datasets consisting of representative eukaryote proteins from all available major eukaryotic supergroups, and viral proteins excluding over-represented Human Immuno-deficiency Virus-1, were clustered into protein families using Diamond BLASTp and Markov Clustering. Hidden-Markov models (HMMs) of clustered protein families were used to search a dataset of prokaryote proteins, limited to 150-bacterial top hits per genus, and the resulting eukaryote, viral, and prokaryote protein families were merged and re-clustered. For a full overview of HGT dataset curation, including software versions and settings used, see Irwin et al.^98^. For a phylogeny of *CLP* in eukaryotes, *Pb* 186b *CLP* and *Homo sapiens* (*Hs*) *SI-CLP* sequences were used to conduct clade-specific searches of the NCBI protein database (gathering threshold: e-01, December 2023), including all available major eukaryotic supergroups and avoiding over-representation from metazoan and plant taxa. These were added to an *SI-CLP* orthologue dataset (OrthoDB) to give a final dataset of 1,445 candidate homologous amino acid sequences.

For all datasets, protein sequences were aligned using MAFFT v7.508^99^ and masked using TrimAL v1.4.15^100^ allowing for no gaps. Each alignment was manually inspected, and highly divergent or identical sequence paralogs from the same genomic source were manually removed. Phylogenies were generated using IQ-TREE v2.2.0.3^101^ with 1,000 ultra-fast bootstraps. Models for each tree were calculated using IQ-TREE’s ModelFinder implementation according to Bayesian inference criterion (BIC) and are detailed in the relevant figure legends. Bootstrap methods and models chosen for tree generation are listed in the respective figure legends.

### RNAi feeding

A full protocol for RNAi feeding and imaging in *Pb* is available at: dx.doi.org<u>/10.17504/protocols.io.8epv5jzm4l1b/v1</u>. Briefly, *Pb* was fed with *E. coli* transformed with an L4440 plasmid construct with paired IPTG-inducible T7 promoters, facilitating targeted gene perturbation through the phagotrophic delivery of complementary double-stranded (ds)RNA. L4440 plasmid constructs (**Supplementary Table S7**) were transformed into *E. coli* HT115 competent cells and grown overnight on LB agar (50 µgmL^-1^ Ampicillin and 12.5 µgmL^-1^ Tetracycline) at 37°C. Positive transformants were picked and grown overnight in LB (50 µgmL^-1^ Ampicillin and 12.5 µgmL^-1^ Tetracycline) at 37°C with shaking (180 rpm). Overnight pre-cultures were back-diluted 1:25 into 15 mL of LB (50 µgmL^-1^ Ampicillin and 12.5 µgmL^-1^ Tetracycline) and incubated for a further 2 hours under the same conditions, until an OD_600_ of between 0.4 and 0.6 was reached. *E. coli* cultures were then supplemented with 0.4 mM IPTG to induce template expression within the L4440 plasmid, and incubated for a further 3 hours under the same conditions. *E. coli* cells were pelleted by centrifugation (3100 x *g* for 2 mins), washed with sterile NCL media, and pelleted once more. *E. coli* cells were then re-suspended in NCL media supplemented with 0.4 mM IPTG, 100 µgmL^-1^ Ampicillin, and 0.8 µgmL^-1^ β-sitosterol, 2% glycerol, and adjusted to a final OD_600_ of 3. *E. coli* cells were split into single-use aliquots and stored at -20°C until feeding^102^.

*Pb* cells were pelleted by gentle centrifugation in a 96-well plate (10 mins at 800 x *g*), taking care not to disturb the cell pellet by leaving 100 µL of supernatant, and re-suspended 2:5 into 150 µL of diluted, induced *E. coli* culture media (to make 250 µL total; final OD 0.1). Feeding was conducted daily for 6 days using frozen bacterial stocks and, where cycloheximide was added alongside feeding, this was added on days 4 and 5 to a final concentration of 50 µgmL^-1^. The orientation of construct position on the microwell plates was altered between experiments, ensuring different columns and rows were used for each construct between experiments (see analysis below).

### Live cell imaging

High-throughput imaging was performed on an ImageXpress Pico Automated Cell Imaging System (Molecular Devices) at 4x magnification, using the Cy5 (absorbance: 630/40 nm; emission: 695/45 nm) channel to capture algal-chlorophyll autofluorescence as described previously^29^. Images were automatically stitched together using the built-in CellReporterXpress image acquisition software, generating a composite tiled image for each well for downstream analysis. Individual cells in each well were masked using the MetaExpress software and the average fluorescence of each object scored. Data for RNAi screening (**Fig. 2a**) and glycan substrate / bacterial feeding (**Fig. 6a, e-g**) experiments were gated by shape [ 0.6 < shape factor < 0.9], size [ area < 10,000] and fluorescence [ average fluorescence intensity < 150] to select for single intact *Pb* cells during cycloheximide treatment.

Average per-host fluorescence data for RNAi screening experiments (Fig. 2a) were analysed using a linear mixed effects model using the lme4^103^ and lmerTest^104^ packages. The model included separate random intercepts accounting for row position, column position, and experimental block to control for batch and/or positional artefacts during imaging. Degrees of freedom to calculate p-values were estimated using the Satterthwaite approximation. Average per-host fluorescence data for glycan substrate / bacterial feeding (**Fig. 6a, e-g**) experiments were analysed using a linear model on log-transformed data (**Fig. 6a, e-g**). All analyses were performed using R version 4.2.1.

### RNA sequencing and differential expression analysis

*Pb* cultures (6x replicates) were subject to RNAi feeding (as above) for 6 days prior to harvesting, including a penultimate 2 days of cycloheximide treatment for the corresponding samples. *Pb* cells (∼50,000 per sample) were strained three times through a 40 μm cell strainer to remove large debris, collected on an 11 μm filter, washed with NCL, and resuspended into 1 mL of TriZol reagent. RNA was extracted using the Zymoprep RNA extraction kit and stored in nuclease-free water at -20°C. RNA was processed using the Illumina NovaSeq 6000 v1.5 workflow (Illumina™) with polyA selection. A 150 bp paired-end library was prepared and resulted in 1,184,917,880 reads total consisting of 177,737,682,000 bp (SRA accession: SRR1251101).

Paired-end reads were trimmed to remove adapters and primers using the TrimGalore v0.6.7-1 wrapper tool of Cutadapt^105^. Trimmed reads were then aligned to our *Pb* 186b Pacbio genome assembly^19^ using BOWTIE2 v2.5.1^106^, and contamination filtered using Samtools v1.7^107^ to keep only host *Pb* mapped sequences (80-90% of all reads). Normalized transcript expression values for each read data set were calculated and cross-sample normalization performed using Salmon v1.10.2^108^. Differential expression analysis was conducted using DESeq2 v1.34.0^109^ to assess change in relative transcript abundance between pairwise data sets.

For four-way analysis of gene expression, pb186bvf_008968-T1 (corresponding to *Pb CLP*) and 11 transcripts (with highly variable or over 75% zero read counts) were removed prior to differential expression analysis to aid in downstream clustering of less differentially expressed transcripts. These sequences were retained in each pair-wise analysis of gene expression. For pairwise and four-way analysis of gene expression, differentially expressed transcripts (|log2 fold change| ≥ 0.5, adjusted p-value < 0.001) were clustered based on normalized read counts. Hierarchical clustering was performed using Pearson correlation and a maximum height cut-off of 80%. Transcript annotation was conducted by manual curation of differentially expressed transcripts using InterProScan.

### Mass spectrometry

For metabolomics following *CLP* RNAi, *Pb* cultures (6x replicates) were subject to RNAi feeding (as above) for 7 days prior to harvesting. *Pb* cells (∼40,000 per sample) were filtered onto 11 µm nylon filters (Merck Millipore NY1104700) to minimise residual bacteria. The cells were washed with water, and the cells resuspended from the filter in 500 µL 80% v/v HPLC-grade methanol. The samples were transferred to 1.5 mL microcentrifuge tubes, snap frozen in liquid nitrogen, and stored at -80°C. Samples were subjected to LC/MS analysis using a derivatised C18 method by the metabolomics facility at the University of Oxford, Department of Chemistry. Data were subjected to sample-specific normalisation based on manual cell counts, followed by median normalisation, log transformation, and Pareto scaling using MetaboAnalyst v5.0^110^ (https://genap.metaboanalyst.ca/). Data were then analysed using a one-way ANOVA, followed by a Fisher’s Least Significant Difference test with false discovery rate (FDR) adjusted p-values (based on the Benjamini–Hochberg procedure^111^).

For metabolomics of *Pb* 186b under differential symbiont load, cultures were first grown at 50 μ mol m^-2^ s^-1^ to increase cell densities, and then split and acclimated to their treatment conditions of low (6 μ mol m^-2^ s^-1^) or moderate-high (50 μ mol m^-2^ s^-1^) light intensity for 3 days. They were then sampled and analyzed as described previously^24^. Briefly, *Pb* cells were filtered on 11 µm nylon filters, washed in Volvic water, and then the cells were disrupted by sonication (20% power for 10 s). The *Pb* fraction was collected by pushing 1 mL of the lysate through a 1.6 µm filter, which caught the intact endosymbiont cells. The samples were analyzed with a Synapt G2-Si with Acquity UPLC, recording in positive mode over a large untargeted mass range (50 – 1000 Da). A 2.1×50mm Acuity UPLC BEH C18 column was used with acetonitrile as the solvent. Masses of interest were investigated using the MarVis-Suite 2.0 software^112^ (http://marvis.gobics.de/), using retention time and mass to compare against KEGG^113^ (https://www.genome.jp/kegg/) and MetaCyc^114^ (https://biocyc.org/) databases.

### Immunofluorescence microscopy and Western Blot of the Pb CLP proteins

*Pb* cells were prepared for immunofluorescence microscopy using the protocol available at: dx.doi.org<u>/10.17504/protocols.io.yxmvm2ozng3p/v1</u>. Primary rabbit polyclonal antibodies for CLP were generated by Thermo Fisher Scientific, using the peptide sequence detailed in **Supplementary Table S7**. Imaging was performed on a ScanR Olympus with a 20X objective (refraction index = 1; emission = 698 [chlorophyll], 515 [Alexa Fluor 488]). Images were collected using cellSens Dimension Software v3.2 and analysed using FIJI ImageJ2 version 2.3.0^115^. In brief, background noise was removed from the z-stack images using the ‘rolling ball background subtraction’ (r=15). To count and analyse the fluorescence of the algae, a smoothing gaussian blur filter was applied, and the z-stacks were combined to create a 2D image. To resolve any overlapping cells, a Watershed segmentation was used after thresholding the image with Otsu method. Processed images were false-coloured to aid visualisation. Cells were manually scored based on whether they exhibited phagosome-like fluorescence (localized to the oral groove and early endosome), lysosome-like fluorescence (localized to small lysosome-like structures clustered around endosymbiotic algae), or both (**Supplementary Table S5**). Algal fluorescence per-host cell and maximum *CLP* fluorescence per host-cell were analysed using a linear model (**Fig. 4c/d**).

For Western Blot, *Pb* cells were first cleaned from debris by straining the culture through a 40 µm cell strainer and washing the cells with autoclaved NCL media. The cells were collected and concentrated via centrifugation (800 x *g* for 10min). Proteins were extracted for 30 min in 2x SDS buffer (55°C) supplemented with a mixture of protease-inhibitor (Roche; Cat.4693116001), 2 µl of RNAse A (Qiagen; Cat. 19101) and 2 µl DNAse I (Fisher Scientific; Cat. EN0521). The detailed protocol for the Western Blot of *Pb* is available at: dx.doi.org<u>/10.17504/protocols.io.ewov1q58kgr2/v1</u>. Imaging was performed on an iBrightCL1000 imager.

### Yeast heterologous expression of Pb putative chitin interacting proteins

*S. cerevisiae* Y16947 was transformed with a p426 GPD plasmid (ATCC 87361) containing a gene expression insert modified with a 5’ mating factor α (MFA) signal peptide sequence, or an empty p426 GPD plasmid control, as described previously^49^.

Transformants were grown for 7 days on SCM-URA at 30°C with shaking at 180 rpm, then centrifuged at 3,200 x *g* for 5 mins and the supernatant collected. Supernatant from each culture was centrifuged at 3,200 x *g* for 40 mins using a Spin-X UF concentrator 10,000 MWCO (Corning) to collect proteins over 10 kDa. Collected protein was washed in PBS and matched to a concentration of 120 μg/mL prior to use. Protein activity assays were conducted using 10 μL of sample (3x replication) on a Chitinase Activity Kit (Sigma-Aldrich #CS0980) following the manufacturer’s instructions, with incubation at 37°C for 3 hours. Comparisons of catalytic function were analyzed using a generalized linear model with quasi-binomial distribution (**Fig. 5d**).

### Symbiont load assessment using flow cytometry

*Pb* cultures to assess symbiont load in **Fig. 6c** and **Fig. S10** were grown under varying light conditions (0, 6, 12, 24 and 50 μ mol m^-2^ s^-1^) with a light-dark (LD) cycle of 14:10 h for seven days. Symbiont load was estimated using a CytoFLEX S flow cytometer (Beckman Coulter Inc., CA, USA) by measuring the intensity of chlorophyll fluorescence for single *Pb* cells (excitation 488nm, emission 690/50nm) and gating cell size using forward side scatter^59^.

### Glycan substrate and bacterial feeding

*Pb* cultures were washed twice with NCL and filtered using a 40 μm cell strainer to remove large debris prior to feeding. For substrate feeding, *Pb* cells (∼150 in 300 μL) in a 96-well plate were supplemented with 0.01 mM, 0.1 mM, or 1 mM of glycan substrate in NCL media (**Supplementary Table S8**). Concentrations of each substrate used are stated in the respective figure panels. *Pb* cultures (6x replicates) were maintained under normal growth conditions at either standard (25 μ mol m^-2^ s^-1^) or high (80 μ mol m^-2^ s^-1^) light intensity for up to 72 hours prior to imaging.

For bacterial feeding, bacterial stocks were grown overnight for 48 hours, washed twice in NCL, and fed to *Pb* cells (∼75-100 in 250 μL) at a final OD_600_ of 0.1. *Pb* cultures (6x replicates) were maintained under normal growth conditions for up to 48 hours prior to imaging (see below).

### Algal growth assays

To determine if *M. conductrix* SAG 241.80 can utilize GlcNAc as a carbon source for heterotrophic growth, growth assays were performed in continuous darkness over a 7-day period in BBM+V media (CCAP) supplemented with 1 mM L-arginine using Biolog Phenotype MicroArrays™ PM1 plates. Populations (3x replication) were initiated at a density of approximately 5 x 106 cells/ml and changes in optical density (OD740) were regularly recorded. Heterotrophic growth on GlcNAc was compared with negative (no carbon source) and positive (α-D-glucose control treatments).

To determine if *M. conductrix* SAG 241.80 can utilize GlcNAc as a nitrogen source, growth assays with 1 mM GlcNAc supplementation were performed over a 10-day period and changes in optical density (OD740) were compared with negative (no nitrogen, -N) and positive control (1 mM L-arginine supplementation) treatments. The algae were nitrogen starved in BBM+V media (CCAP) lacking a nitrogen source (i.e., BBM+V-N) for 24 hours before the growth assays commenced. Populations (6x replication) were then initiated in 96 well plates at a density of approximately 5 x 106 cells/ml in BBM+V containing 100 mM MES and 100 mM MOPS buffer and, where applicable, 1 mM of the nitrogen source to be tested at pH 5.5, 6.5 and 7.5. Plates were incubated at 24°C with a 14/10 h LD cycle for 10 days and OD740 readings were regularly recorded.

