## Supplementary Figures S1-12 for "Immune-like glycan-sensing and horizontally-acquired glycan-processing orchestrate host control in a microbial endosymbiosis"

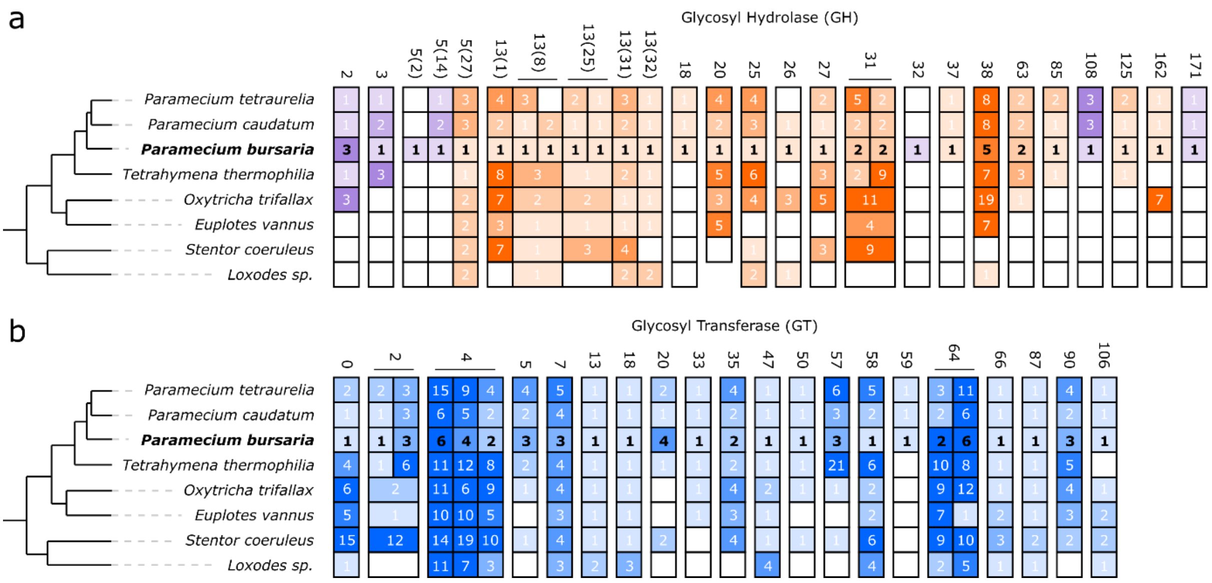


**Supplementary Figure S1**: Coulson plot showing the distribution of (***a***) glycosyl hydrolase (GH) and (***b***) glycosyl transferase (GT)-domain containing genes in *Pb* and representative ciliate species[57]. Genes were identified using a custom CAZY annotation and tree-building pipeline (see **Methods**). Numbers above each column indicate the CAZY family annotation. Orange indicates putative 'ancestral' GH genes; blue indicates putative 'ancestral' GT genes; and purple indicates genes putatively acquired through horizontal gene-transfer (HGT). Shading and numbers indicate the number of copies (in-paralogues) of each gene family identified in the genome, with subdivided boxes (e.g., CAZY annotation 13(8)) indicating *Paramecium* specific gene duplications. Phylogeny schematic based on Irwin et al.[57].

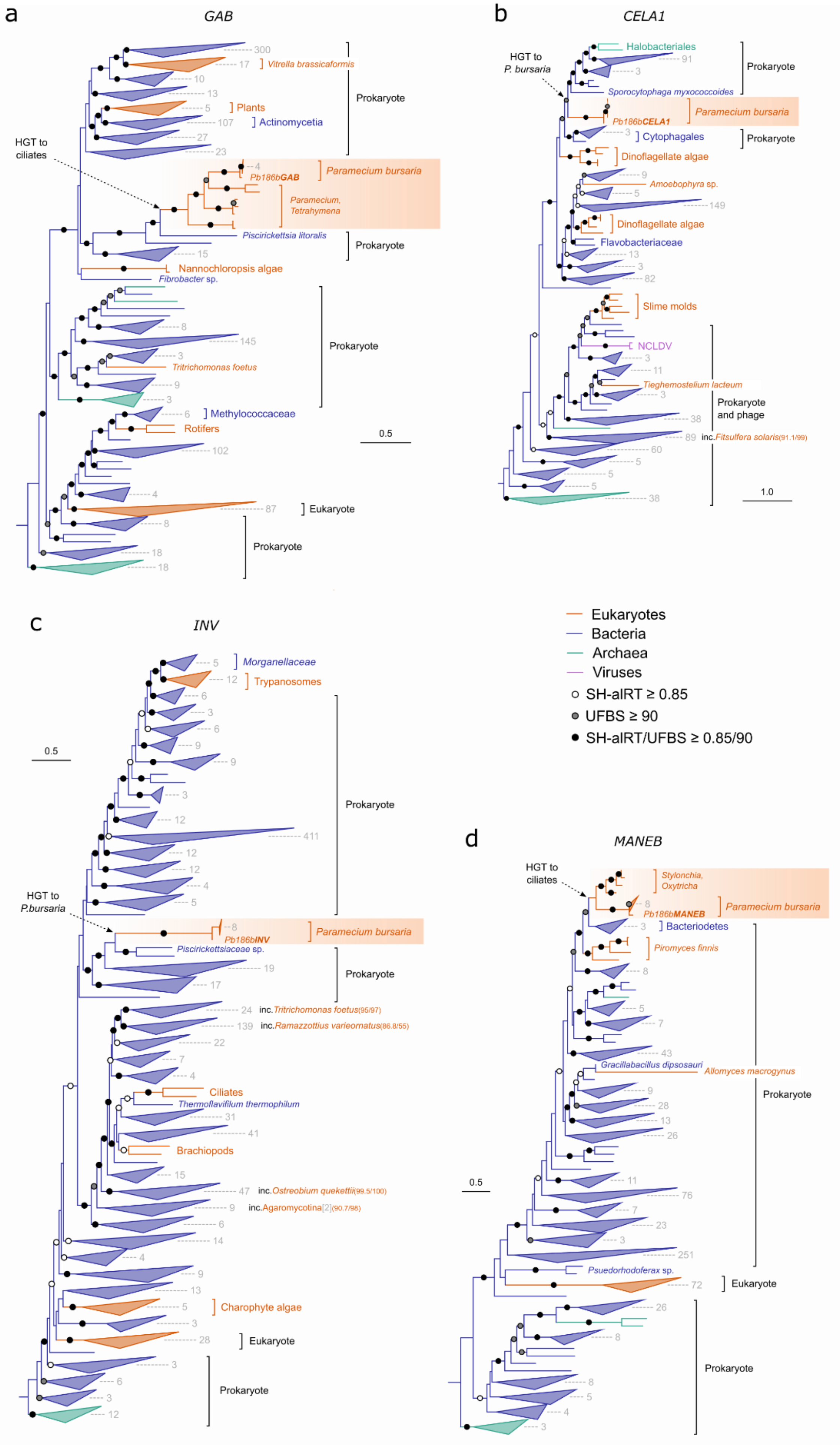
a
GAB
HGT to ciliates
Vitrella brassicaformis
Plants
Actinomycetia
Prokaryote
Paramecium bursaria
Pb186bGAB
Paramecium, Tetrahymena
Piscirickettsia litoralis
Nannochloropsis algae
Fibrobacter sp.
Tritrichomonas foetus
Methylococcaceae
Rotifers
Eukaryote
0.5
b
CELA1
HGT to P. bursaria
Halobacteriales
Sporocytophaga myxococcoides
Pb186bCELA1
Cytophagales
Dinoflagellate algae
Amoebophyra sp.
Flavobacteriaceae
Slime molds
NCLDV
Tieghemostelium lacteum
Prokaryote and phage
inc. Fitsulfera solaris(91.1/99)
1.0
c
INV
Morganellaceae
Trypanosomes
HGT to P.bursaria
Pb186bINV
Piscirickettsiaceae sp.
inc. Tritrichomonas foetus(95/97)
inc. Ramazzottius varieornatus(86.8/55)
Ciliates
Thermoflavifilum thermophilum
Brachiopods
inc. Ostreobium quekettii(99.5/100)
inc. Agaromycotina[2](90.7/98)
Charophyte algae
Eukaryotes
Bacteria
Archaea
Viruses
SH-aLRT ≥ 0.85
UFBS ≥ 90
SH-aLRT/UFBS ≥ 0.85/90
d
MANEB
Stylonchia, Oxytricha
Pb186bMANEB
Bacteriodetes
Piromyces finnis
Gracillabacillus dipsosauri
Allomyces macrogynus
Psuedorhodoferax sp.

**Supplementary Figure S2:** Maximum likelihood phylogenies of *Pb β-glucosidase* (*GAB*; ***a***), *Cellulase (CELA1*; ***b***), *β-fructofuranosidase* (*INV*; ***c***), and *Endo 1,4-β-mannosidase* (*MANEB*; ***d***) genes. Topologies of these trees provide strong support for prokaryote-to-eukaryote HGT. Trees were generated in IQ-TREE using Q.pfam+R10 (***a***), LG+R9 (***b***), Q.pfam+I+R10 (***c***), or Q.pfam+R9 (***d***) substitution models selected by Model Finder, and statistical support assessed using SH-aLRT and ultra-fast bootstraps (UFBS; *n* = 1000).

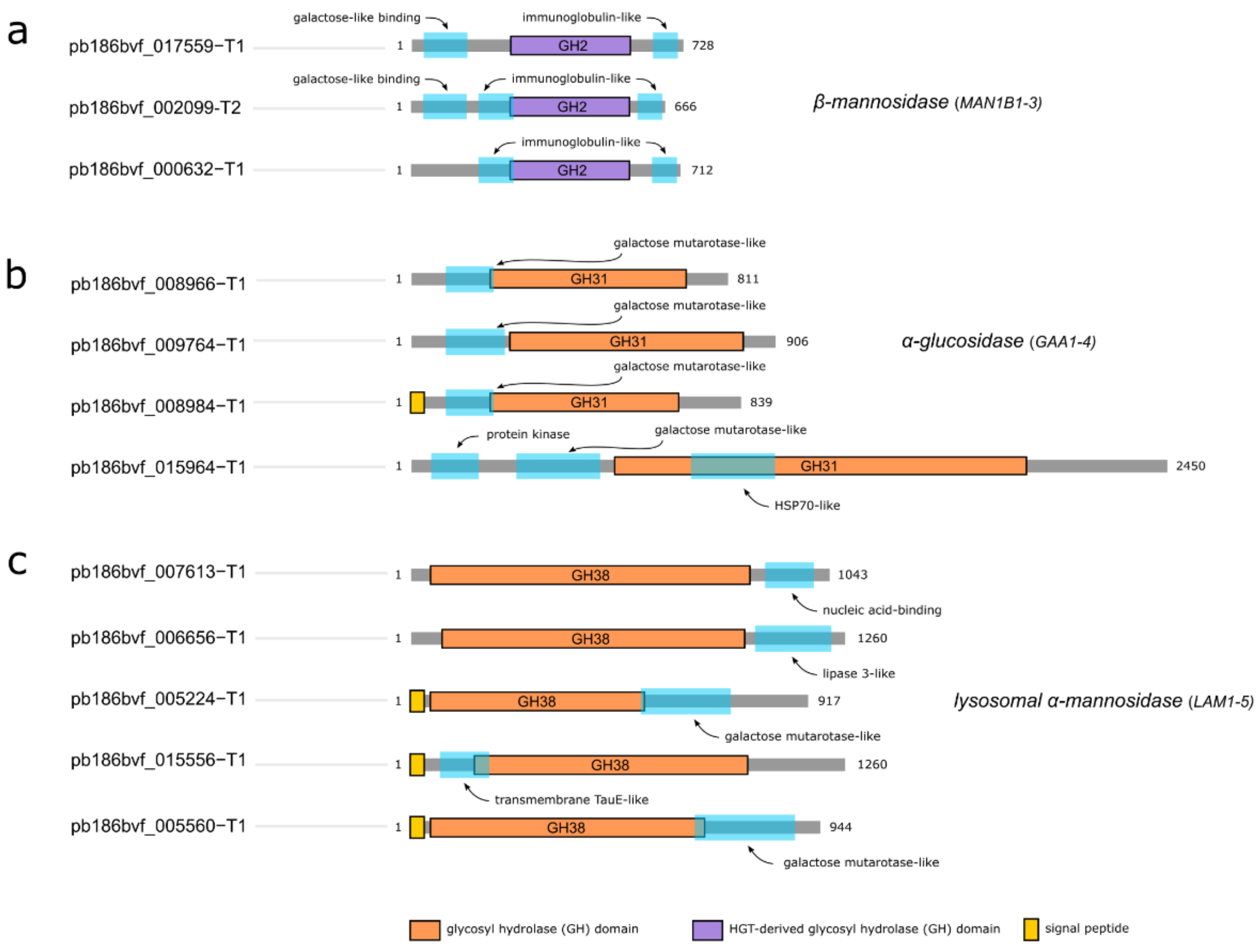


**Supplementary Figure S3**: Schematic illustration showing functional domain architecture of individual *Pb β-mannosidase* (***a***), *α-glucoside* (***b***) and *lysosomal α-mannosidase* (***c***) transcripts used in chimeric RNAi constructs (see **Fig. 2b**). GH, glycosyl hydrolase; additional domains are highlighted in cyan.

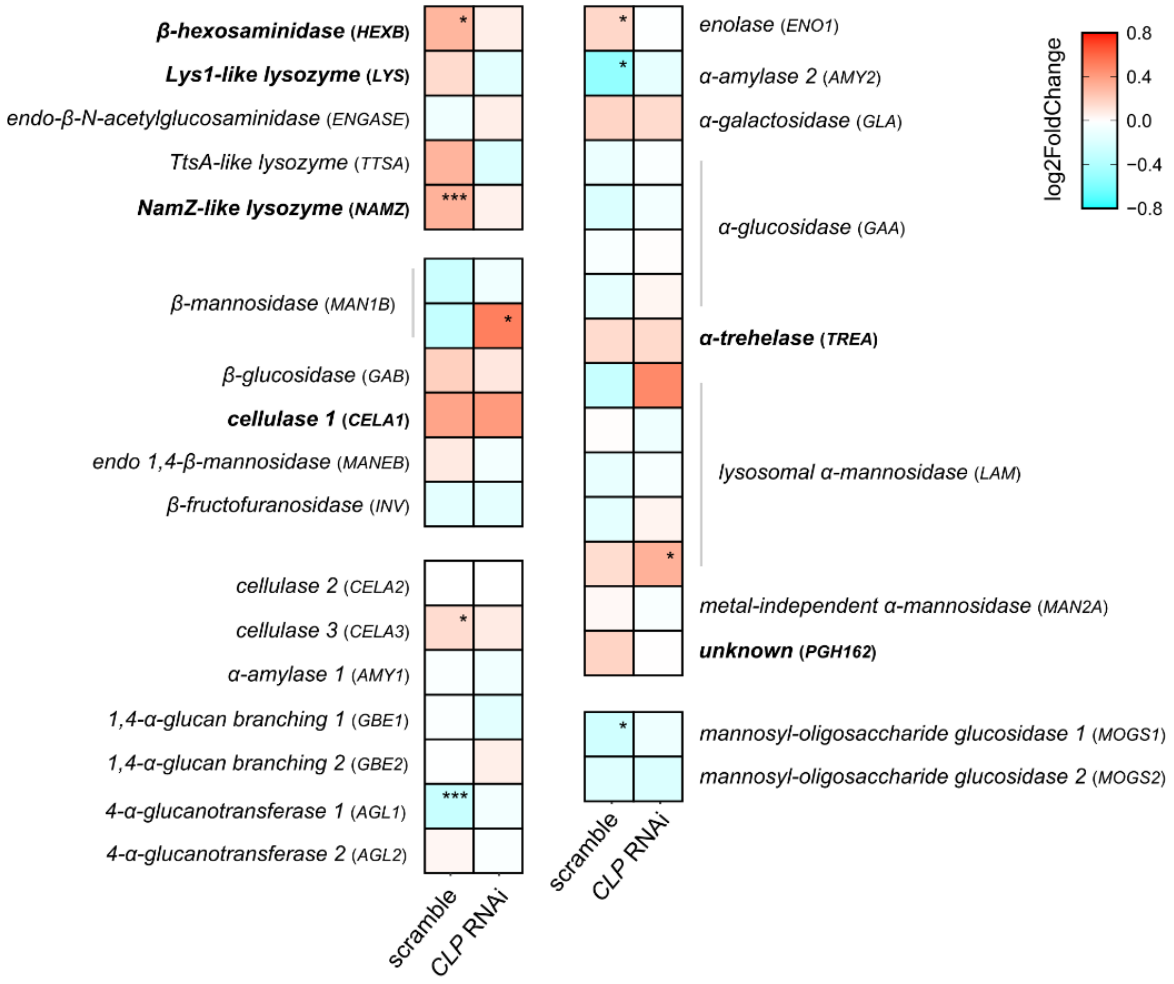


**Supplementary Figure S4:** Differential expression of GH transcripts during cycloheximide treatment to induce endosymbiont break-down, compared to untreated controls. Cultures were simultaneously exposed to non-hit 'scramble' or *CLP* dsRNA. Note the perturbation of *HEXB*, *LYS1* and *NAMZ* gene expression in response to *CLP* RNAi. Significance calculated as *p ≤ 0.05, and ***p ≤ 0.001 using an adjusted p value (see also **Supplementary Table S2**).

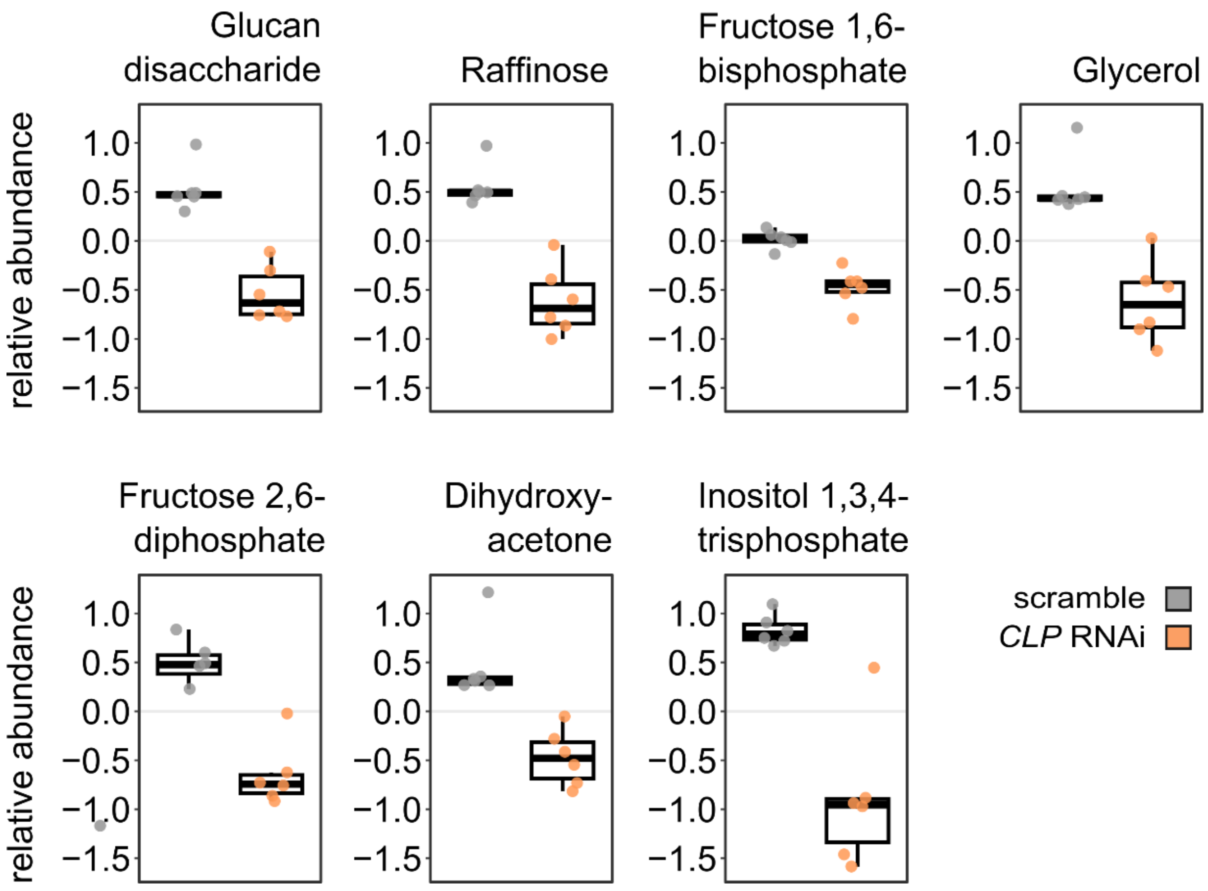


**Supplementary Figure S5**: Relative abundance of significantly altered metabolites detected during *CLP* RNAi. All annotated metabolites with an FDR-corrected p-value <0.05 are shown.

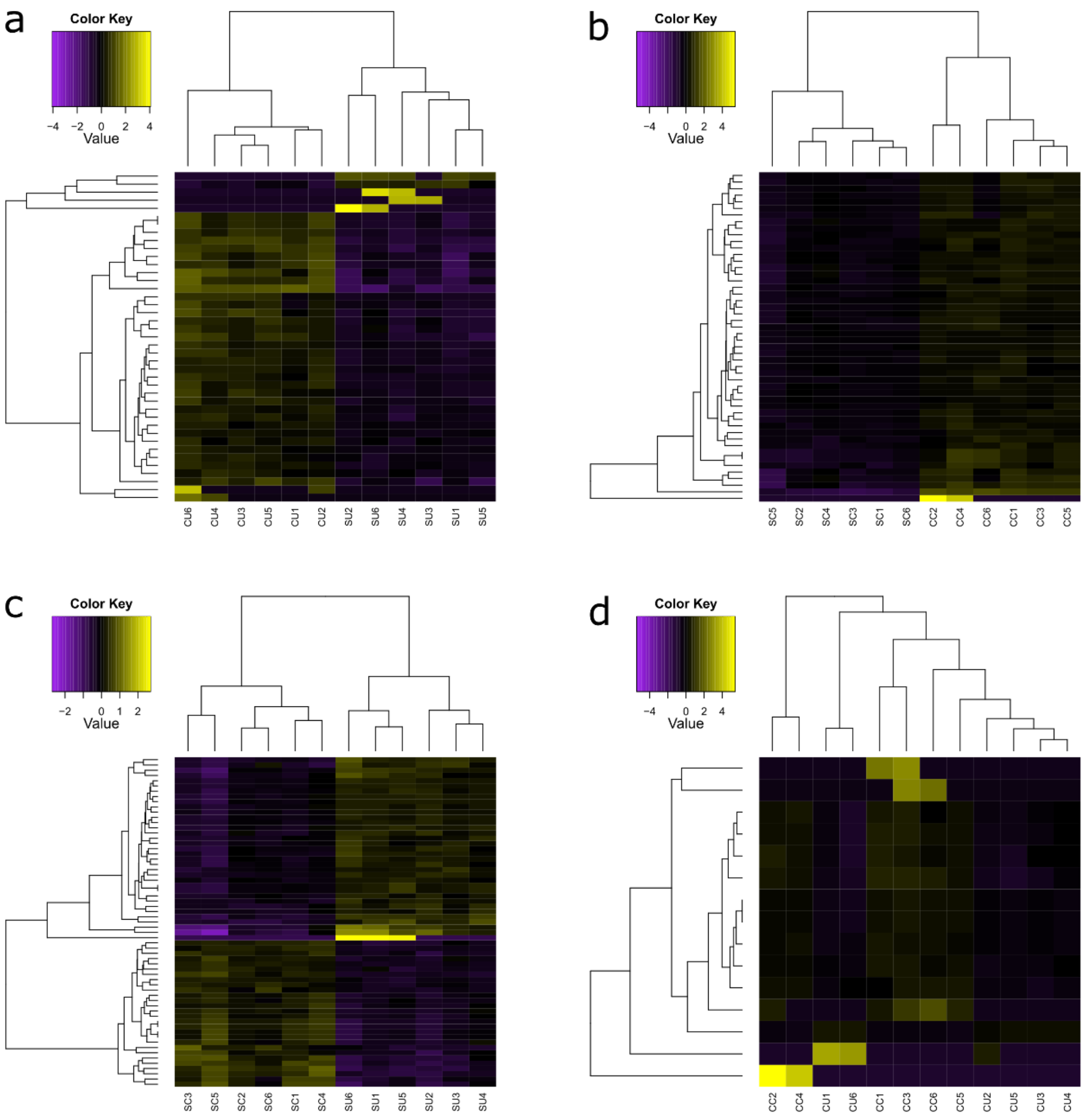


**Supplementary Figure S6**: Differential expression analysis to assess the effect of *CLP* RNAi on genes with altered expression during cycloheximide treatment. Hierarchical clustering of differentially expressed transcripts ([log2 fold change] ≥ 1, adjusted p value < 0.001) in each pairwise comparison; ***a***, scramble vs *CLP* RNAi [no cycloheximide], **b**, scramble vs *CLP* RNAi [+ cycloheximide treatment], ***c***, cycloheximide vs no cycloheximide [scramble], and ***d***, cycloheximide vs no cycloheximide treatment [*CLP* RNAi]. Note the reduction in differentially expressed transcripts upon cycloheximide treatment during *CLP* RNAi in ***c*** vs ***d***.

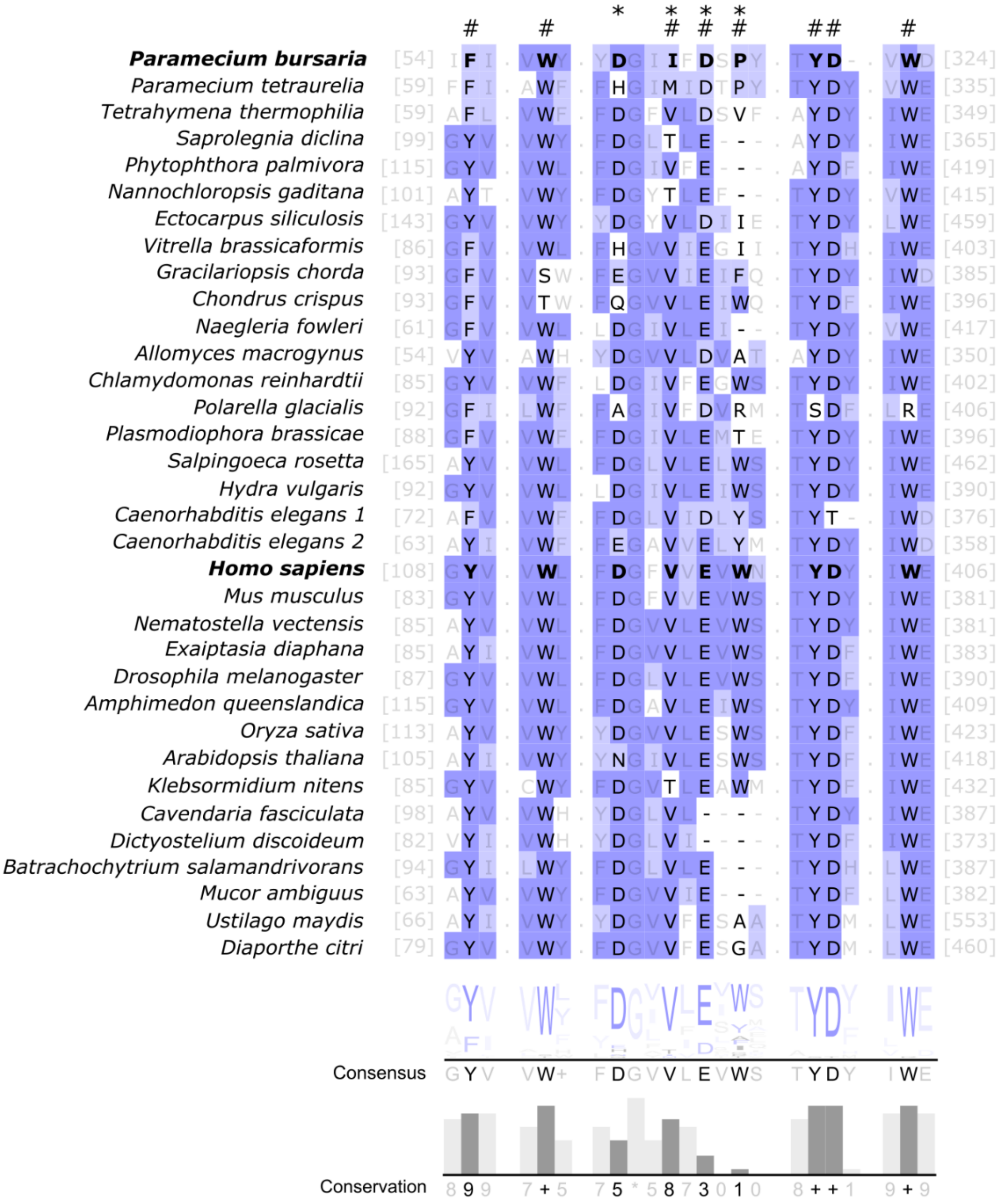


**Supplementary Figure S7**: Trimmed amino acid alignment showing conserved functional domain residues of *CLP* across eukaryotes including characterised binding (#) and cleavage (*) site amino acids. Amino acid consensus and conservation values are shown below. Purple shading indicates BLOSUM62 scores indicating evolutionary divergence of individual sites.

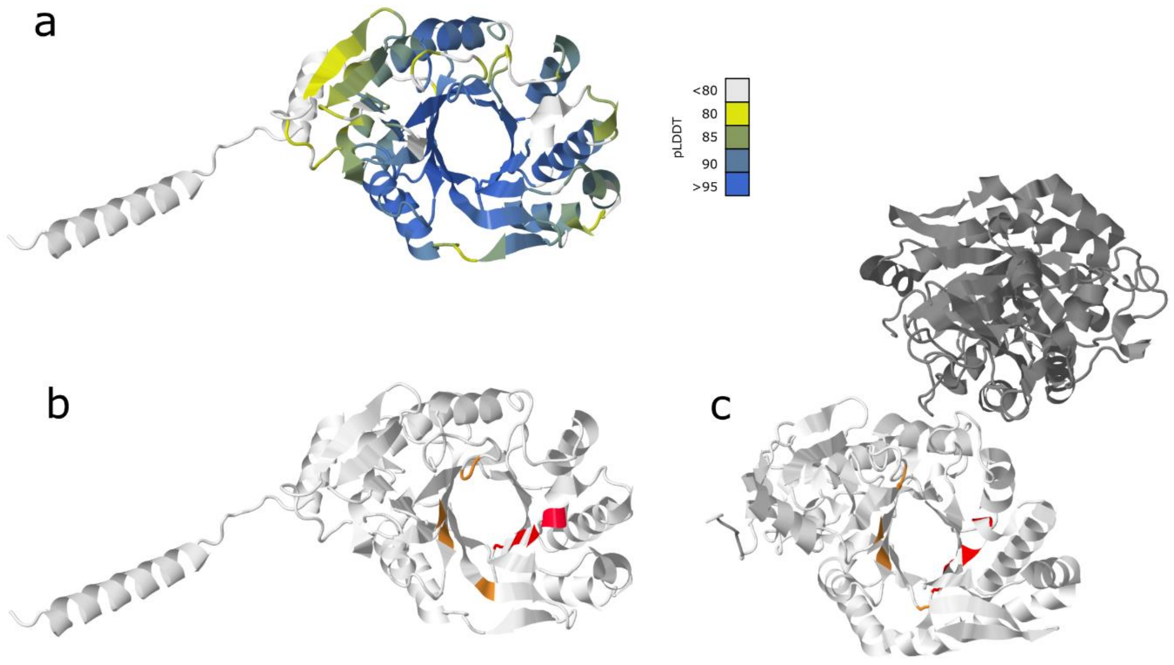


**Supplementary Figure S8:** ***a***, Putative 3D structure of *Pb CLP* predicted using AlphaFold3. Shading indicates pLDDT model prediction confidence. ***b,*** Putative 3D structure of *Pb CLP* compared with ***c***, the crystal structure of *Homo sapiens SI-CLP*. Characterised binding (orange) and cleavage (red) sites are highlighted. Note that *H. sapiens SI-CLP* is predicted to form a dimer.

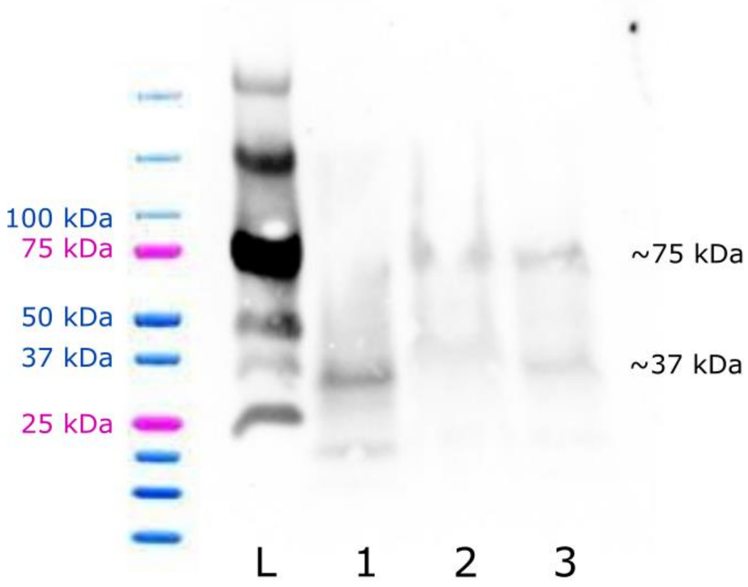


**Supplementary Figure S9**: Western blot of *Pb CLP* from two independent sample preps. Cells were harvested via filtration followed by centrifugation (1) and 1:2 dilution (2), or via two-step centrifugation (3). Ladder (L) and size representation are shown. The predicted molecular weight of *Pb CLP* is 40.24 kDa) Note that two bands are visible in each sample corresponding approximately to 37 kDa and 75kDa, suggesting that *Pb CLP* may also form a dimer.

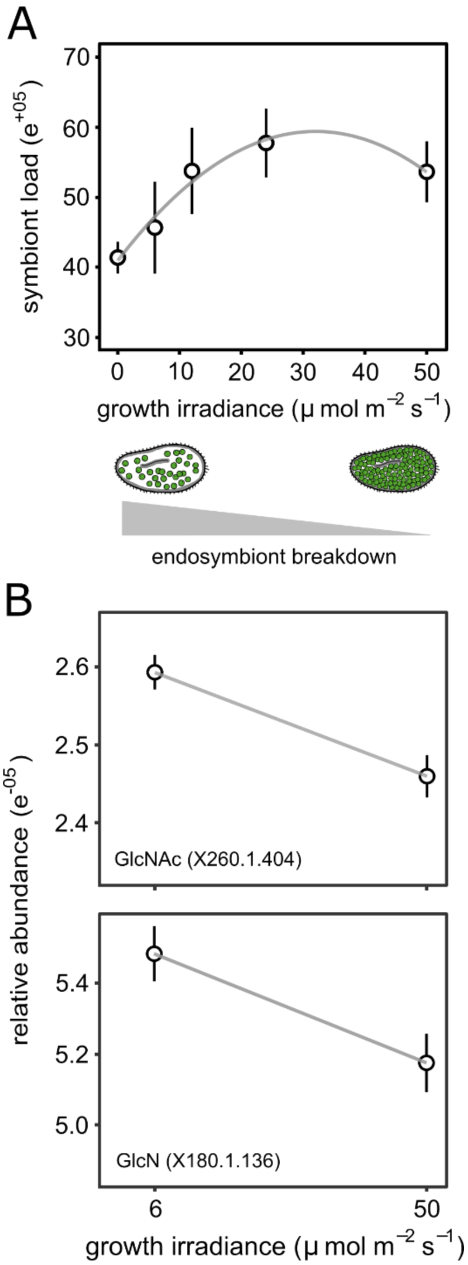


**Supplementary Figure S10:** Symbiont load (***a***) in *Pb* cultures grown for 7 days under 0, 6, 10, 25, or 50 μ mol $m^{-2}$ $s^{-1}$ light irradiance. Data are plotted as mean ± SE of 3 biological replicates. (***b***) Relative GlcNAc and GlcN metabolite abundance at two light levels for two populations of cells. Data are plotted as mean ± SE of 5 biological replicates.

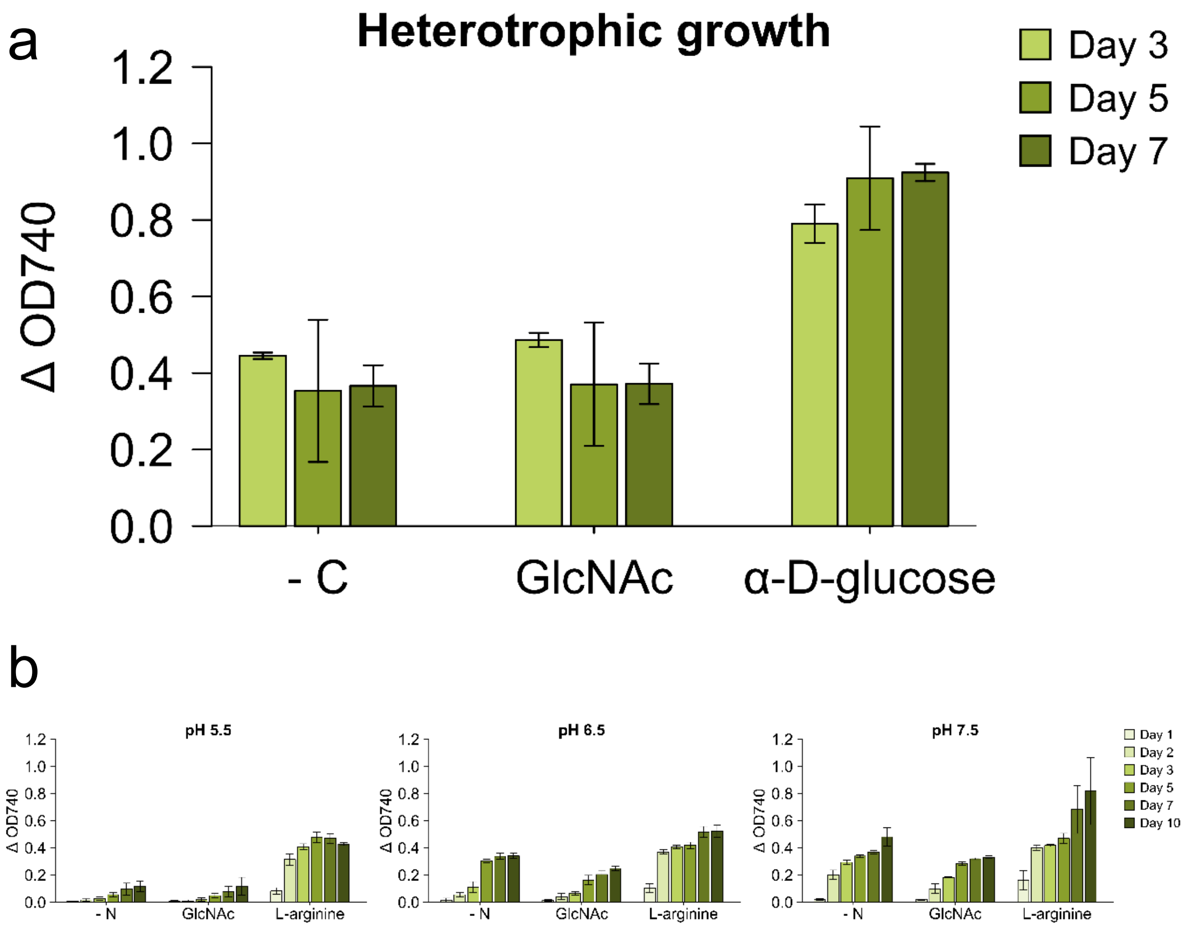


**Supplementary Figure S11:** ***M. conductrix*** **SAG 241.80 cannot utilise GlcNAc as a carbon source for heterotrophic growth or as a nitrogen source.** Heterotrophic growth assays were conducted in constant darkness using Biolog Phenotype MicroArrays™ PM1 plates with BBM+V media (pH ~ 7) and 1 mM L-arginine supplementation (***a***). Growth with GlcNAc was compared with negative controls lacking a carbon source (-C) and positive controls supplemented with α-D-glucose. Utility of potential nitrogen sources was assessed by comparing growth with BBM+V media lacking a nitrogen source (-N) or with 1 mM GlcNAc or 1 mM L-arginine at pH 5.5, 6.5 and 7.5 (***b***). For all experiments, growth was determined based on changes in optical density (OD740) over time relative to baseline values (Day 0)

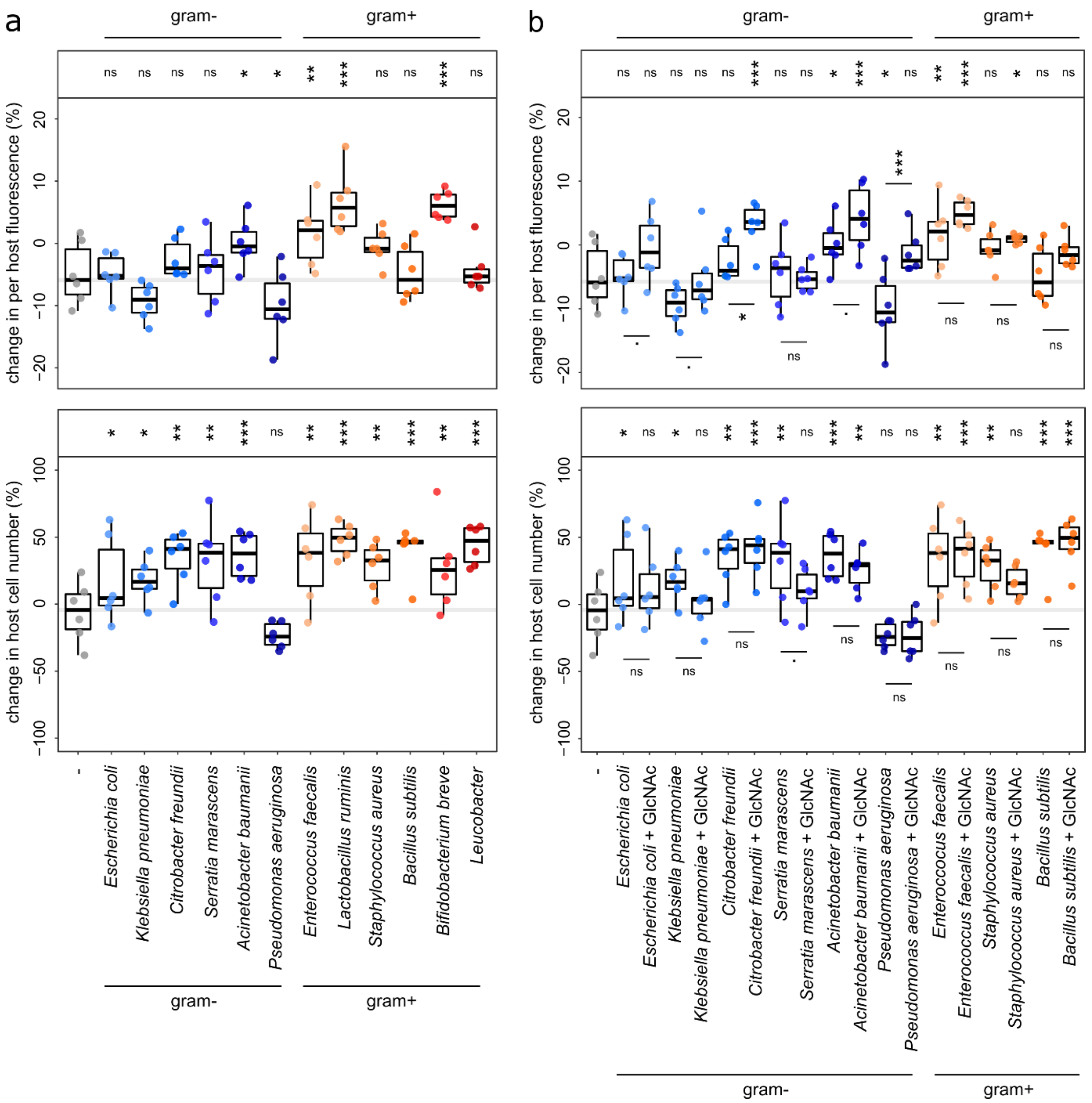


**Supplementary Figure S12**: Percentage change in per host fluorescence, and host cell number, in *Pb* cultures following bacterial feeding (***a***) or bacterial feeding plus exposure to 1mM GlcNAc (***b***) for 24 hours. Bacteria were heat-killed and fed to *Pb* at a final $OD_{600}$ of 0.1. Boxplot data are represented as max, upper quartile (Q3), mean, lower quartile (Q1) and min values, and individual data points for each biological replicate are shown. Significance calculated as *p ≤ 0.05, **p ≤ 0.01, ***p ≤ 0.001, and ns = no significance, using a linear model on log-transformed data. Significance values above each plot denote individual comparison to the unfed control (-), and significance values within the plot denote pairwise comparisons between indicated variables. Note that the greatest increase in per host fluorescence resulted from exposure to gram positive (gram+) bacterial strains, and that co-exposure with 1mM GlcNAc increased per host fluorescence during exposure to gram negative (gram-) bacteria.
