## Supplementary Table S7 for "Immune-like glycan-sensing and horizontally-acquired glycan-processing orchestrate host control in a microbial endosymbiosis"

Scramble	| pBJ078
AAGCTTGTACGGGGGGACGCGATCCCCTTAGCCGGCGCAGATTTTTGTCGCTCACATGACTGCCGCGAACGACGGTTCCCGTAATGGACCAACCACCAGGCCGGATGCTCGAGGCTGACATTTGAGCGCCACGCGCCAAATGTAAATCGGGCAAGCCTGCAG

U2AF	| pb_epi2
CTGCAGATAGTTTGATGGTTATTCCTTTCCATTAACTTTCTTAACTTATGCAGTATCTCCTTAAACTTTACAAATTTTTAATCTCTTATCACCAATTTAAAGATTATTTAATGCTTTAAGAGCTTTTACTGTAGATGATGGGTTTTCGTATTCAAAAAAACAATATCCTTTTGAAATATCACTATTGCTCGTCGTGTCTTTGACAAGATTGAAATATTTTGTAATGCCAAAAGTTTGAATTAGTTTTAAAACATCATCATCTTTCAAATAAGTTGGTAATCCTCCTAAATATAATCTCGTATCATCTAATTTCAATTCCGTCACTTATTCATTGGGTACAGGATTTAAAATTCTCTCTAAAAATTTTCTAGGCCTTTCTATCTTTAATTTGTGATTGACAAATTCAACTTGATTTAGAGGCAGAAGGGTCCTTTTGGCTTCTATAGAACTACATTCTAACACAACCCATGATTTGGTTGCTCCATACTCTATGCTTTTAATTGGCTAAGCTT

CLP	| pBJ033
AAGCTTGGATATTAATATACTTTAGATTACGCGTAAAAAATAGATTATGTGTCACCTGTTTGGTATAATATAGAAAAAGATTTATAACTAAATGGCAATAATGAGAATCCTATATGGATTAATGATATAAAAAAGAAGAATCCTAATATAAAGATCCTGCAG

HEXB	| pBJ034
AAGCTTAATTTAATTATGACGCAGCTTTTGCTACTGTTATGCAAAATATCCTTCGTAATATCTCTGATGCCTCAACCCAGAGTGATGAAAAATGGTACTGTGACTAGAAAATTTGAACCATGTTAAACAATTTACGTATCAGAATAAATGTTTCCACTGCAG

LYS	| pBJ035
AAGCTTAAAAGGCTCCTTCTGGTATTGCTTGTCGTCCTTCCATGCTTAGCTGTCAAGGGAGTTGACCTAAGTTAAGCATTTAATAACTTTGCCTGCATAAAGAGCAATGGATATTCGTTTGCCATCGTGAGAGGTTATATGTCCTATGGAGCAGTGCTGCAG

ENGASE	| pBJ053
AAGCTTTCAAATACTTCATCTACTACTGCACCTAAAAGAAGTATTTCGTGTGATTAACTAACTACAAAACTGAAGAGTCTCAAGTCAGCATACGATAGAAAATAATATTTGAAATTTATAGCTAGATAGAGAGAAATCTAACGCGATCAGGAAGAGCTGCAG

TTSA	| pBJ054
AAGCTTTTTAATATTATGTTTTAGAATTGGCTATCAGGATAATCAAATACAAGTAATTTTTCGTACTCTGATCCATTTTCTCCTACCAATATTGAAGACTTTGAAGATTGTTTAAAATATTTAGAATAAAAGTAACCTAATGATATTTAATAAATACTGCAG

NAMZ	| pBJ067
AAGCTTCTTAATCACAAATCCAACAGGAGTGGATGATGATCTCACAATGATAACTGATAATATAGGTTAATATTGCAATATTCGGAAATTCTTTTCGCCTGAACATGGATTAAGAGGCGATAAATAAGCAGGAGTTGCTGTTGATGACTATATTGACTGCAG

MAN1B_chimera	| pBJ060
AAGCTTCAGATATGTTTATGCCAAGAGCACTTAAGAATCCTCAGGTTTATGTTGATATATTTTTAAATTCTTTGGCAGCAGGATATAATGGTTTGAGAATTTGGGGAGGAGGATAGTTTGAATATGATATATTCTATGAATTGGCAGACCAACATGAACATTACCTTTATATGTTTATTGTCTATCATTAACAACATAAACATTAATTGAACTATTATCAGTTAAATTAGCGAACATGGCTGGATCTTTGTGACTCTTTTTAGCAGCATAATGGCCTCCTTTCCAACAATTGTAAAAATCAACTGTTAATGGGTTAAGATTAATCAAAAATACTTGAGAAATAATGGATTTACAGAAGCATAAATGTATTGAATATTACAAATGCTGAAATTATATTTGAAGGAATAGACACTATTGCATCAATTTATGTCAACAATCAGAAAGTGATAGATGCTACTGCAG

GAB	| pBJ022
AAGCTTATTTTAATGATCAATATTGCAGCTGGTGGGAAAATATTTTCACAGCACTACGATCTCGCTTAAGAAATAGTTAATTCTATGTCCTTGGATTAAAAAATTGGATAGACCCTATAAATTGATTATGTCGATATCACAAGAGGACAAACTGATCTGCAG

CELA1	| pBJ061
AAGCTTTACATTATTGTGCATTGGTGTAGCCATGAGCTAAACTCCAGTTTAGAGATGGGGAAGATTGAGAGTTCAAGGAAATCAATTAGTGTCTGAAAGCGGATAGGCAGTTCAACTTAAAGGTATGTCCACTCATGGTCCTCAATGGTTTGCTAACTGCAG

MANEB	| pBJ063
AAGCTTCATAAGCATTATACCTAAGATTACAGAGCAAATAAGGCACACATATAATTAGTGGATAATTTGACCATTACAGCGACATTCAATCAATAACTGGTAAAGAGCCTGCCATTAGAGGATTTGACTTATAAAACTATTCCCCTCACAATCCTTCTGCAG

INV	| pBJ043
AAGCTTTTATTTATTTAATTAGCATTATGTATTTAAGAGAAAGCAATAAATTTTGAAGCCTCAAATTCTAAAAGAGAAATCATACTATTGTATTAGAAAAAGATAGATTAAGCAATAAACCTAGCTATTAATGCTGAATAATAACCTCTTCATAACCTGCAG

CELA2	| pBJ024
AAGCTTTTAATTGCATTCGTCATATTTGGAGTCTACGCACATGTGTAGACTGACATCAGATATGGAAAAGTTCAGAGCAAAGGAGTAAATCTAGGTGGCTGGTTAGTTGCTGAGCATTGGATGACTTCAGATTCTGTCATTTGGGCTGGGGTTTCTCTGCAG

CELA3	| pBJ025
AAGCTTATGTTCTTGCTAGGCCTAATAGTATTGGCAAGCTCTTTCCGAATTGACAGAGATGATTAATTTATTGTTGATGAAGATGGATTAAGAAGAGTTTATCATGGTGTTAATGTTGTCTATAAGGTTGCTCCCTTCTATCCACCTATTTCAGAGCTGCAG

AMY1	| pBJ026
AAGCTTACTAAGGAGGAGTGGAAAAGTCGAGTAGTCTACTAATTGTTAACTGATAGGTTTGCTACCAGTAAAGGAGCATCAACTAGTTGTAATTTAGGTAATTACTGTGGAGGAGACTATAAGGGAATGATTGAACACTTGGATTATGTTAAAAACCTGCAG

GBE1	| pBJ062
AAGCTTGAAGAAGAAATTGATGAGAAAACCGGACCACCAATAATTATATTTCCAAAAACACCTGTGAAAATACATGAAACTCTTATTAGCGATTTTGCTGATTGTGATGAACCAAAAAGATCAATGGGGCTCATATTGGGGAGAAAATAGGTGGATCTGCAG

GBE2	| pBJ028
AAGCTTGATAGGGAAGCATATAAACCATACGAAAGAAAGGTAGCGAAAAAATCGTCTTAATTTTACAAAATGGCATTAGAAGAGAAGTTAGAATTCCACTAAAAATATGAAAAAGATACTAATCAATTGGAGTTATTTTAACAAACACCAGAATTACTGCAG

AGL1	| pBJ029
AAGCTTATTAATCTAAATCCTGAAATTCAAATCGAAGAAAGAGTAGTTTACTTCTATTAGGGTGATACTTGTCGCTTCATCCTATAAAACAAACTTTAGTATCAGAATCCTAAGATTAAAATATAAAAAGAGGAAATCAAATTGGCACTTGGTGAGCTGCAG

AGL2	| pBJ030
AAGCTTAACAATCATGAATAAAAGTTTTCATCTCTATCAACGAGAACCTTAGACATAATTGATGAAATAGTGACTGTAAGGAGTGAAAATTACACCTTAGAATAATCCAGATTATTAACTGCCTTCACTACTCCAATGAAACCAATTAAAAATAAACTGCAG

ENO1	| pBJ031
AAGCTTGTTGAATATTTGGAATGTTTCAAGTGTGGGTAGCAAGTCGATTAAATACTTGCTCTCTAGTGTGACCATAATCTTTGTTTACAATGTGCAGCTCGAGCTGTATAAAAATGGTAAGGTAGTTAATAGCCATTGATTTGTTCCATATGTGGACTGCAG

AMY2	| pBJ032
AAGCTTTATCTTGTCTTAGCACTAGCTGCTGCAGCATTTATTGTCGTGCTCTATGCACCCAATAATACCAATAACCTTCGTGAAACATGCAATAACTTCGATAATGAGAATGCATGTTAGAATGGTCAATAGACAGAGAACCCAACTACATGGACTGAGCTC

GLA	| pBJ037
AAGCTTTCAGTTTATGCAATTGATAATGGTCTTGGTCTCACTCCACCCATGGGATGGAACCCATGGAATAAGTATGGTTGTGAGATTAGCGAAGACTTGATTAAGTAAGCAGCAGATAGGCTTGTATCCACAGGTTTGGCAGCAGCTGGATATATCCTGCAG

GAA_chimera	| pBJ066
AAGCTTGCTTATTATTATGCAACATTGATTGAATCAAATGTTACATTTAAATAATACAGCCTAAGGCCAAAAGAAGTGAAGCAGTTGAGGGTCAATGTCTCAACCATTCAATTAGAGGAAGAATTAATTTTTGAACTAATTGAGCATGGATCGTAGTTCAGATTATAAAAATGGATATTCACATAATTGGTAATTAGATATGACAAATAAATTGGTGCAGCTGTTGGTAGTTTAATATAACCATTATGGTGGTTTGAACCTAATGATCCAATAGCAATATAATATGAAGAGAGAGAATTTATATTTATATAATTATTACTTGAATTCACAAGGATCAGTATTTAGGCTAGAAAGATTAGATGGAACTGTTCTATTTGATCTGAAATCCACTGATGTATTATCATTACTTTATCATGAATTACATGTATAAAAAGAAAGCAATTACATTTGGGGGCTAGGAGAAAGAGCCCAATCATCCAGATTTCGGAATACTTCTAGAAGGGACTGTAATAGTTACTCAAGAAACTCCAACTCAATTGGACATCAGAATAGAAGATGCAAAGAATGAATTGTTCAGATTTCCTGACATTGAGCCATTTAAATCAGTCTTCAATGACTGCAG

TREA	| pBJ044
AAGCTTAAATTAGCTTAGGAAATAAATAAATTATGGCTGGGATTATCTAAAAGATTCACATATGACGAATCAAAATCATCTTACATCAGTTGCCCTCATCCATTTTTTGTGCCAGGAGGAAGATTTAGAGAATTTTACTATTGGGATACACTCTGGCTGCAG

LAM_chimera	| pBJ050
AAGCTTCGGCACAACCTATTAGTTGGACAAAGAAACCATTTGGATTATAAGGGTTCTAACAATCTTATTAGAGTACATTTGATTCCGCATTCTCATGATGATGTAGGATGGCTAAAAACGTATGAGGAATATTATTATGGACTCAACAATCGGGTTTTATTATTAGTGTTACTTTCCTTAACTGCTTCTAAGTTTTTAGGATAAGTGAGTATCGTCAAAAGTGACATAAGAGAAGTGCATCTTATACCTCATTCCCATGATGATGTAGGTTGGTTGAATACTGTGGAAGAAATTTATTAAAGAGGAAGAGTTGCAGCTATCTAACTCTACATAGGAAAATAGTAGGAGTATTTAAACATTAATATGATTATAATTTTATTAATATTTAAAAATATATTAGCCCACAAATAATATAATGAAACAATCGCTTAATAATTGATAGCCATGAGCTTTGCATTATTGATATATTTTGTATACAGTCAAACTGTAAATGAGAGAGTAGCTGAATGTAACCCTAAAATAGTTTGCGAAACAGTACAATAAACCTTAAATTTAAATCTACATTTTACATTGCATTCTCATTTAGATGCGTTTTGGCTGTAAACATTGTTATTTTCATTGGTGGCCGCAAACCTCAGATTTGGTGACAAGGAACAATTGGAAAAGCACCATAAACATTTCCCTTTAGTAAATGAATCTCTGACAGTTCATTTAATACCACATTCACATGATGATGTGGGTTGGCTTAAAACTGTTCTGCAG

MAN2A	| pBJ055
AAGCTTTCCAAAATCAAGGATTAGAACTTTAGTATAATATTTACAAACACATTCCCTAATACTCTTGATACAACAGTTATGAGTTTTACTGTTGATTAATTTGGAAATCCAGATTCATACATAATTACGGGAGATATTGATGCAATGTGGCTTAGACTGCAG

PGH162	| pBJ056
AAGCTTCTAGTTGTGATAGTGACATCTTAGAGTGATTGCAGATTTGCATTCAAATATTCATAAGAAGACCTAAAAACTAAACCTGAAGTCTTATCAGAATTTTTAAAAACAGTTGTTTAGTGGGAATCAAAATTTGCTAAAACTATTGGCATAGACCTGCAG

Pb_CLP_epitope	| OBS-29:47
ETFKELLDKELANEIDLTK
