## Supplementary Table S8 for "Immune-like glycan-sensing and horizontally-acquired glycan-processing orchestrate host control in a microbial endosymbiosis"

**Supplementary Table S8: Compounds used in this study**

| **Compound** | **Source** |
| --- | --- |
| D-glucose | Sigma-Aldrich G8270 |
| D-glucosamine hydrochloride | Alfa Aesar A15532.18 |
| D-glucosamine 6-phosphate | Sigma-Aldrich G5509 |
| N-acetyl-D-glucosamine | Sigma-Aldrich A8625 |
| N-acetyl-D-glucosamine-6-phosphate disodium salt | Biosynth MA06746 |
| Diacetyl-chitobiose (Chitin_2_) | Megazyme O-CHI2 |
| Triacetyl-chitotriose (Chitin_3_) | Megazyme O-CHI3 |
| Triacetyl-chitotriose (Chitin_4_) | Megazyme O-CHI4 |
| Pentaacetyl-chitopentaose (Chitin_5_) | Megazyme O-CHI5 |
| β-Sitosterol | Sigma-Aldrich 567152 |
